# Improved reconstruction of transcripts and coding sequences from RNA-seq data

**DOI:** 10.1101/2025.02.27.640589

**Authors:** Jan Grau, Jens Keilwagen

**Affiliations:** Institute of Computer Science, Martin Luther University Halle-Wittenberg, Halle, Germany; Institute for Biosafety in Plant Biotechnology, Julius Kühn-Institut, Quedlinburg, Germany

## Abstract

**Motivation:** Annotation of genes and transcripts is an important requirement for understanding the information that is encoded in newly sequenced genomes. One source of information suited for this purpose are RNA-seq data mapped to the respective genome sequence. RNA-seq-based approaches for transcript reconstruction generate transcript models from these data by combining regions of contiguous coverage (exons) and split read mappings (introns). Understanding phenotypes as a consequence of proteins encoded in a genome further requires the annotation of coding sequences within transcript models.

**Results:** We present GeMoRNA, a novel approach for transcript reconstruction from RNA-seq data that combines a combinatorial enumeration of candidate transcripts with heuristics for splitting candidate transcripts in regions of contiguous coverage and subsequent likelihood-based quantification. We benchmark GeMoRNA against the previous approaches Cufflinks, Scallop and StringTie using a large collection of public RNA-seq data for seven species. For the majority of species, we observe an improved prediction performance of GeMoRNA, especially on the level of coding sequences and for species with dense genomes. We combine GeMoRNA with the homology-based approach GeMoMa to yield a re-annotation of two recently sequenced genomes of *Nicotiana benthamiana* lab strains.

**Availability and implementation:** The source code of GeMoRNA is available from GitHub at https://github.com/Jstacs/Jstacs/tree/master/projects/gemorna. A binary version of GeMoRNA is available from https://www.jstacs.de/index.php/GeMoRNA. The annotation files for the *N. benthamiana* lab strains are available from zenodo at https://doi.org/10.5281/zenodo.14901380.

## INTRODUCTION

Gene prediction is one of the longest-standing problems in bioinformatics, and pioneering approaches date back to the 1990s (1, 2). Accurate predictions of genes, transcripts and transcript variants are essential for virtually any downstream tasks in genomics and transcriptomics, including the construction of pangenomes (3), phylogenomics (4), the detection of orthologs (5) or the analysis of differentially expressed genes or transcripts (6). Since many phenotypes are expressed via the proteins encoded by (protein-coding) genes (7), the annotation of coding sequences (CDSs) within transcripts is an important additional requirement for functional genomics (8).

Methods for computational gene prediction can be roughly categorized into *ab initio*, homology-based and expression-based approaches. *Ab initio* gene prediction uses mathematical or statistical models to represent typical sequence signals of, for instance, coding exons or splice sites to predict transcript models in newly sequenced genomes based on the genomic sequence alone. The most popular instance of a (mostly) *ab initio* approach today is Augustus (9–11), while the recently developed Tiberius builds upon specialized neural network layers and has been reported to further improve prediction accuracy (12).

In homology-based gene prediction, protein sequences or transcript models from well-annotated reference species are mapped to the newly sequenced genome based on sequence homology on the level of (translated) protein sequence. Splicing in eukaryotes can be considered by performing spliced alignments in approaches like miniprot (13) or Spaln (14), or by including the conservation of intron positions as a feature like in GeneMapper (15) or GeMoMa (16). Finally, expression-based approaches start from experimental evidence of expressed transcripts, which initially have been cDNA or EST sequences (17), but today are mostly next- or third-generation RNA sequencing data. Hence, we will refer to such methods as RNA-seq-based in the following. One pioneering approach for gene prediction, also termed transcript reconstruction in this context, from RNA-seq data has been Cufflinks (18). More recent alternatives include Scallop (19) and StringTie (20), where the latter is currently the most popular choice according to citations.

Mixtures and combinations of these prototypical approaches have been implemented in methods and pipelines like MAKER2 (21) or BRAKER3 (22), and many tools allow for the inclusion of hints from at least one additional source of evidence. For instance, Augustus and GeMoMa both make use of additional RNA-seq-based evidence (23–25).

Each of the prototypical approaches has specific limitations. Purely *ab initio* gene prediction is prone to false positive exon predictions and errors in general (22, 26). RNA-seq-based approaches may only predict genes and transcript variants that are sufficiently expressed and, hence, depend on the diversity of the sequencing libraries considered. By design, homology-based predictions are limited to those genes/proteins that are present in the reference and may, hence, miss orphan, i.e., species-specific genes.

The latter limitation also applies to our homology-based prediction method GeMoMa. Hence, we aimed at developing an RNA-seq-based prediction method that may complement the GeMoMa predictions to also yield predictions for genes that are not present in the reference species. Since the homology-based prediction of GeMoMa is intrinsically limited to protein-coding genes, this approach should also focus on protein-coding genes and include an internal prediction of coding sequences (CDSs) within transcript models. While non-coding RNAs play important roles in an organism, protein-coding genes are still a common starting point for elucidating the functions encoded in a genome (8). In addition, the joint prediction using a homology-based and an RNA-seq-based approach allows for identifying high-confidence transcript models with double evidence.

This novel approach, termed GeMoRNA, differs from previous approaches in two central aspects. First, we generate candidate transcripts combinatorially from the exons and introns with RNA-seq evidence, which are subsequently filtered by likelihood-based quantification. Second, we apply several heuristics that split overlapping candidate transcripts candidates while taking CDS predictions into account. We compare the prediction performance of GeMoRNA with the previous approaches Cufflinks, Scallop and StringTie in benchmark studies considering diverse RNA-seq libraries for seven species, namely the three plant species *Arabidopsis thaliana, Oryza sativa* and *Solanum lycopersicum*, the three animal species *Caenorhabditis elegans, Drosophila melanogaster* and *Mus musculus*, and the yeast *Saccharomyces cerevisiae*. We further illustrate how the combination of homology-based (GeMoMa) and RNA-seq-based (GeMoRNA) predictions may be used to balance sensitivity and precision. Finally, we apply the combined pipeline of homology-based and RNA-seq-based (GeMoRNA) prediction to re-annotate two recently *Nicotiana benthamiana* genomes.

## METHODS

### GeMoRNA algorithm

The GeMoRNA algorithms starts from a set of reads mapped to the respective reference genome. Mapped reads are then used to build a base-resolution *read graph*, which is further processed into a *splicing graph*, which roughly represents exons and introns. Based on the splicing graph, candidate transcripts are enumerated, which are further tested for possible splits, quantified, and filtered to yield the final prediction. In Figure 1, we illustrate the general workflow of the algorithm, while each of its steps are described in detail in the following.

**Fig. 1.**
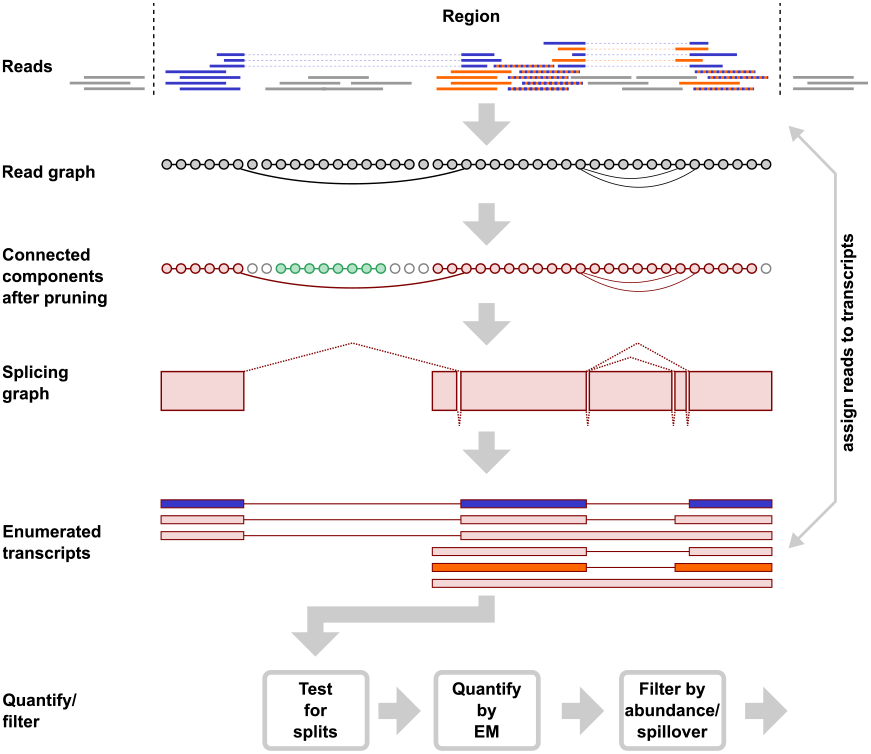
Schematic representation of the GeMoRNA algorithm. First, a nucleotideresolution read graph is constructed from the mapped reads, which is decomposed into connected components and converted to a splicing graph. Based on the splicing graph, candidate transcript models are enumerated combinatorially and tested for possible splits. Reads are assigned to compatibility classes of transcript models as indicated by the coloring of two example transcripts and corresponding reads in blue, orange or striped in both colours. Transcript abundance is quantified by an EM algorithm on the compatibility classes of reads, and finally filtered and reported.

### Defining genomic regions

The input of the GeMoRNA algorithm are mapped reads in (coordinate sorted) BAM format together with the strand orientation of the sequenced library. Reads are filtered for mapping quality (default: 40). To reduce computational complexity, we first partition the genomic sequence into *regions* with contiguous coverage by mapped reads. In this step, we count a genomic position covered, if either a read base has been mapped to this position or if it is contained in the split region (as indicated by “N” stretches in the cigar string) of a mapped read. Small gaps in the coverage profile (default: 50 bp) may be closed by adding *dummy* reads, which are disregarded in later quantification steps. In case of a strand-specific library, regions are defined for each strand separately. Regions with less than 10 reads in total are discarded. If the coverage on one strand is 50-fold larger than the coverage of the other strand and the Jaccard coefficient of the sets of covered positions in the two regions exceeds 0.5, both regions are merged, retaining the strand orientation of the regions with the larger coverage.

If the length of the resulting region exceeds a user-defined threshold (default: 750 kbp), the region is split into subregions at the positions with the lowest coverage. This is repeated until the length of each sub-region is below the defined threshold. The final region or sub-regions, respectively, are used to build the read graph in the following step.

### Generating and processing the read graph

The read graph is generated separately for each genomic region. In the read graph, each genomic position within the region is represented by a node. Two nodes are connected by an edge if two adjacent bases of a read have been mapped to the respective position. This may apply to nodes of the read graph that represent directly adjacent genomic positions if these are contained in a read stretch that has been mapped contiguously, but also to nodes representing more distant genomic positions in case of split reads (cf. Figure 1, Read graph). In the latter case, edges are only considered if they connect nodes that are more distant than a user defined threshold (default: 10 bp) as these may later be considered as putative introns of transcript models. For each node and each edge, we further record the set of supporting reads.

Edges are subsequently pruned by absolute and relative coverage according to user-defined thresholds. After pruning, the read graph is decomposed into connected components to reduce computational complexity in the following steps.

### Generating the splicing graph

Each connected component of the read graph is further processed into a splicing graph. In the splicing graph, stretches of connected nodes without alternative edges that represent contiguous genomic positions are merged to *contiguous regions* (cf. Figure 1, Splicing graph, red boxes). In case of alternative edges from one node to multiple alternative nodes of the read graph, the respective contiguous regions are connected by edges. This may include edges between contiguous regions directly adjacent in genomic coordinates, but also to non-neighboring or more distant contiguous regions. The latter edges will later be considered as putative introns. Both contiguous regions and edges in the splicing graph, inherit supporting reads from the respective nodes and edges of the previous read graph.

### Enumerating candidate transcripts

Based on the splicing graph, we combinatorially enumerate possible transcript models. Conceptually, we start from each of the contiguous regions that are connected to other contiguous regions only at their right end, i.e. that do not have incoming edges reading the splicing graph from left to right. From one contiguous region, we follow outgoing (right) edges to the next contiguous region one by one, where each alternative edge taken results in the generation of an additional, nascent transcript model. This depth-first procedure is repeated, combinatorially generating nascent transcript models in each step, until we reach a contiguous region without outgoing (right) edge. In the enumerated transcript models, directly adjacent contiguous regions are merged into exons, while edges between these are considered as introns. All transcript models generated in this manner are reported as putative transcripts for the following steps. In some cases, this procedure may result in a combinatorial explosion of transcript models. Hence, if the number of enumerated transcripts exceeds 1000, the procedure is stopped before completion and only transcripts enumerated up to this point are considered. To reduce the loss of putatively relevant transcript models, outgoing edges are considered in the order of their coverage and further coverage-based heuristics are applied to decide if an edge is followed in this case.

### Splitting candidate transcripts

If two neighboring true transcripts are located close to each other on the genome, it may frequently happen that the coverage profiles of both transcripts overlap. With the previously described procedure such true transcripts may share a contiguous region in the splicing graph (as these are defined by contiguous coverage without alternative edges) and, subsequently are represented by a joint transcript model. To counteract this issue, we implemented multiple splitting heuristics that are applied to each of the enumerated transcript models.

These split heuristics include rules for the preservation of coding sequences (CDS). To this end, we apply a simple “longest ORF” CDS prediction, where the spliced sequence according to a transcript model is conceptually translated in all three reading frames, and the longest stretch between a start codon and the next in-frame stop codon is considered as CDS. The same CDS prediction is also applied when reporting final transcript models in GeMoRNA.

A first split heuristic is based on the coverage profile within the candidate transcript model. Here, we search for pronounced “dips” in the coverage profile, i.e. local coverage minima (Supplementary Figure S1), or a pronounced coverage imbalance, where one part of the transcript model has substantially less coverage than another part (Supplementary Figure S2). Specifically, we compute the average coverage in sliding windows of 50 bp (small) and 1000 bp (large) and search for small windows with less than 20% of the coverage of the less-covered, neighboring large window. In addition, we search for position, where the coverage in the preceding large window is at least 4-fold greater/less than in the subsequent large window. Position fulfilling either of these criteria are tested for a possible split. A split is performed only if the original transcript model has at least one intron and the two split candidate transcripts still each contain a proper CDS prediction with a user-defined minimum number of amino acids (default: 70). In case of a split, the two resulting new transcript models may be further split repeating the same procedure.

A second split heuristic is applied only in case of non-strandspecific libraries, where overlapping transcripts may occur in head-to-head or tail-to-tail configuration. Here, we consider the orientation of splice sites within the enumerated transcript (Supplementary Figure S3). The orientation of splice sites may be determined from the canonical donor (GT/GC) and acceptor (AG) di-nucleotides at the first and last positions of putative introns. If we identify an exon in the transcript model that separates a contiguous stretch of introns indicating one orientation from a contiguous stretch of introns indicating the opposite orientation, this exon is considered for a split following the same logic as for the first heuristic but without the requirement of a CDS prediction in both split transcript models. Since we cannot determine an accurate split position from splice sites alone, the exon at the border between the two regions with differing orientation is included in both split transcript models.

A third heuristic is considered only if neither of the two previous heuristics had an effect. In that case, the CDS prediction of the original transcript model is taken into account, and we test if the remainder of the transcript (i.e. 5’ UTR or 3’ UTR according to the original CDS prediction) may harbor another CDS of sufficient length. In addition, we require that both resulting split transcript models contain at least two introns. The latter heuristic shall reduce the number of false-positive splits in long UTRs with declining coverage. If these criteria are met, the original transcript is split into two transcript models.

### Assigning reads to transcripts

For subsequent quantification, reads need to be assigned to candidate transcript models. Specifically, we assign reads to *compatibility classes* based mon their concordance to one or multiple transcript models, which is inspired by the definition of equivalence classes in kallisto (27). A read is compatible with a transcript model if all of its mapped bases fall into the transcript’s exons, and all split regions in the mapping match its introns. By this definition, a read may be compatible with multiple candidate transcript models as illustrated in Figure 1.

We define for read *r* and candidate transcript model *t*

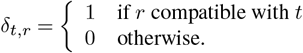

Since contiguous regions and edges in the splicing graph and, hence, exons and introns of the transcript model store lists of supporting reads, the assignment can be solved efficiently by set operations (union, set difference) on those (sorted) lists.

### Quantifying transcript abundance

Quantification of candidate transcript abundance is performed using a likelihood-based approach, again inspired by kallisto (27). Let *R* denote the set of reads that have been assigned to any of the candidate transcripts that originated from the current connected component of the read graph. Let *T* denote the set of those candidate transcripts. Let denote *w*_*t*_ the *a priori* weight of candidate transcript *t*, and *v*_*r*_ the weight of read *r*, chosen according to heuristics described in Supplementary Methods. The likelihood of the set of reads given the abundance vector α of the candidate transcripts is proportional to

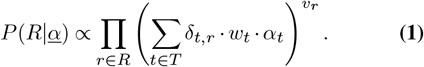

Transcript abundances –_*t*_ are determined by maximizing the likelihood (equation 1) with respect to the vector – using the expectation maximization (EM) algorithm.

### Filtering transcripts

Based on the transcript abundances –_*t*_ obtained in the previous step, candidate transcripts are filtered by absolute and relative abundance. Specifically, candidate transcripts are ordered according the their abundance, and candidate transcripts are added to the output set until

a. the absolute abundance falls below a user-defined threshold (default: 20),
b. the (relative) abundance falls below a user-defined threshold (default: 0.05),
c. the already added transcripts explain a user-defined fraction of the total abundance (default: 0.9), or
d. the fraction 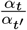 of abundance relative to the next candidate transcript *t*^*′*^ in sorted order is larger than a userdefined threshold (default: 20).

In case of strand-specific libraries, a small fraction of reads may also occur on the opposite strand of the original transcripts, and such “spillover”, single-exon transcripts are filtered by a heuristic based on local coverage. Multi-exon candidate transcripts are also filtered if their strand orientation according to their donor and acceptor splice sites does not match the strand orientation according to the library, and single-exon candidate transcripts are filtered if the theoretical abundance of their CDS (i.e. abundance multiplied by fraction of CDS relative to total length) falls below the threshold on absolute abundance (criterion (a) above).

### Handling of long-read data

For handling long-read (e.g., Pacific Biosciences or Oxford Nanopore Technologies) mapping data, a few, minor adaptations of the described algorithm are required, since current long reads may contain mismatches and insertions/deletions relative to the reference genome at a higher frequency than short (e.g., Illumina) reads, and typically span multiple introns.

First, the requirement of at least 10 reads per region is lifted. Second, edges are pruned from the read graph, if their number of supporting reads is less than a fifth of the number of supporting reads of an alternative edge that shares one node with the edge in question while the pair of second nodes has a distance of at most 5. Third, the weighting of reads in the quantification process is adapted as described in Supplementary Methods.

### Tuning heuristics

Heuristics have been derived and tuned by systematic assessment of prediction performance and manual inspection of critical cases (e.g., missing transcripts, fusion transcripts) based on the following libraries (ENA/SRA accessions): *A. thaliana*: ERR1886195, ERR2245559, DRR09053; *D. melanogaster*: SRR3659025; *M. musculus*: ERR2983488; *O. sativa*: SRR1785721. In addition, longread (PacBio SMRT) data for *A. thaliana* (SRR12350726) have been used for testing the application of GeMoRNA to long reads. None of these data sets have been used to evaluate prediction performance in the benchmark studies.

### Benchmark data

GeMoRNA has been benchmarked on short-read (Illumina) data for *A. thaliana, O. sativa, S. lycopersicum, C. elegans, D. melanogaster, M. musculus* and yeast (*S. cerevisiae*). Genome sequences and annotations of all species have been obtained from the sources listed in Supplementary Table A. For each species, 40 paired-end libraries were selected at random from those present in the European Nucleotide Archive (ENA, https://www.ebi.ac.uk/ena/browser/home) that have been sequenced using Illumina HiSeq or NovaSeq instruments. Due to the focus on protein-coding genes, libraries have been stratified for library selection: 85% of the libraries were sequenced after poly-A enrichment (field `library_selection` cDNA, PolyA or Oligo-dT), while 15% where sequenced after rRNA depletion (Inverse rRNA (selection)). In addition, we aimed at a considerable fraction of strand-specific and non-strand-specific libraries, which is not represented by a standardized field in ENA. Hence, strand-specificity of libraries was determined by custom scripts after mapping the reads to the respective reference genome as detailed below. For *S. cerevisiae*, the initially sampled libraries did not contain a sufficient number of non-strand-specific libraries. For this reason, the number of sampled libraries was increased to 60, yielding 9 non-strand-specific libraries. A list of all sampled libraries (ENA accessions) is provided as Supplementary Table B, where a few libraries failed processing for different reasons (*D. melanogaster*: 2; *O. sativa*: 1; *S. cerevisiae*: 3) (cf. Supplementary Figure S4).

### Benchmarking procedure

Sampled libraries were mapped to the respective reference genome (Supplementary Table A) using STAR 2.7.10b with parameters `--alignIntronMax` and `--alignMatesGapMax` set to 100, 000 for *M. musculus* and *D. melanogaster*, and to 20, 000 for the remaining species to avoid spurious mappings with very long splits that sometimes occur in STAR mappings. Further non-standard parameters were `--runThreadN 4` `--outSAMtype BAM SortedByCoordinate`. Finally, we set `--outSAMstrandField intronMotif` to allow for gene prediction using cufflinks from non-strand-specific libraries.

After mapping, strand specificity of each library was determined using a custom script based on featureCounts (28) using the respective reference annotation (Supplementary Table A). If the number of reads assigned by featureCounts using one strand orientation was at least 2-fold the number for the other strand, the library was reported as FR_FIRST_STRAND (following cufflinks nomenclature) or `FR_SECOND_STRAND`, respectively. If the ratio between both parameters was less than 2, the library was reported as `FR_UNSTRANDED`.

For benchmarking purposes, GeMoRNA, StringTie v2.2.1, cufflinks v2.2.1 and Scallop v0.10.5 were applied to each of the mapped libraries separately.

GeMoRNA was run with default parameters except for parameter `s` set to the key for the respective strand specificity of the library. StringTie (20, 29, 30) was run with default parameters except for `-p 6`, and the parameters for strand specificity set to `-rf` in case of FR_FIRST_STRAND, `-fr` for `FR_SECOND_STRAND` in case of strand-specific libraries. Cufflinks (18) was run with default parameters except for `-p 6`, and `-library-type` set to the key for the respective strand specificity of the library. Scallop (19) was run with default parameters except for `-library_type` set to `first` in case of FR_FIRST_STRAND and `second` for `FR_SECOND_STRAND` in case of strand-specific libraries. The output of GeMoRNA, StringTie, Cufflinks and Scallop was compared against the respective reference annotation (Supplementary Table A) using gffcompare v0.12.6 (31). The gffcompare output was analyzed primarily on the “Transcript” level and is intended to provide a measure of prediction accuracy on the level of transcript models, irrespective of CDS annotations.

On the level of CDS, the output of all four tools was further compared against reference using the “Analyzer” tool of GeMoMa (16, 24, 25) with default parameters. Since StringTie, Cufflinks and Scallop do not perform CDS prediction internally, we followed two alternative strategies: first, we made the simplistic CDS prediction of GeMoRNA applicable to external GFF/GTF files in a separate tool, and applied this tool to the output of StringTie, Cufflinks and Scallop. Second, we applied TransDecoder v5.7.0 to the output of these three tools (cf. Supplementary Methods), since the simplistic prediction built into GeMoRNA might unnecessarily reduce the prediction performance of the alternative tools. We did not apply TransDecoder to GeMoRNA output, since the CDS predictions of TransDecoder might contradict those that have been used by GeMoRNA-internal heuristics when splitting candidate transcripts.

### Benchmarking long-read data

In addition, we benchmarked GeMoRNA on long-read (PacBio, ONT) data. GeMoRNA and StringTie provide a dedicated long-read mode, while Scallop-LR (32) requires PacBio-specific header files and has not been updated for five years. Hence, we limited the comparison to GeMoRNA and StringTie in this case.

Benchmark data have been obtained from ENA for the same seven species as for the Illumina short-read data. Data sets were sampled from those available at ENA with at least 100, 000 reads to yield 5 data sets per instrument of the PacBio Sequel and Sequel II, and the ONT GridION, MinION and PromethION instruments. If less than 5 data sets were available for an instrument, further data sets were selected from the remaining at random. For *O. sativa* this resulted in 15 and for *S. lycopersicum* in 13 data sets, while for the remaining species the desired number of 25 data sets could be obtained. A list of all sampled libraries (ENA accessions) is provided as Supplementary Table C.

Strand specificity of the libraries was determined in analogy to the Illumina short-read data, but with additional flag `-L` for the featureCounts call.

StringTie was run on the long-read data sets with additional parameter `-L` and GeMoRNA was run with additional parameters `lr=true mrpg=5 mrpt=5`, where the first indicates the long-read mode of GeMoRNA, while the other two parameter lower the number of supporting reads that are required to report a gene or transcript.

The remainder of the benchmarking procedure (gffcompare, CDS prediction, Analyzer) was completed in analogy to previous short-read data.

### Homology-based prediction using GeMoMa

Since GeMoRNA shall complement the homology-based predictions of GeMoMa, we additionally used GeMoMa to predict transcript models for all seven species, considering existing transcript models from three closely related species as reference. Conceptually, this scenario is not typical for a homology-based prediction, because the seven well-annotated species considered in this manuscript would rather be used as a reference when annotating newly sequenced genomes, and may have also been used to annotate those considered as a reference here.

The species and genome versions used as reference for predicting transcript models in the genomes of *A. thaliana, O. sativa, S. lycopersicum, C. elegans, D. melanogaster, M. musculus* and *S. cerevisiae* based on sequence homology are listed in Supplementary Table D.

We use the *Extract RNA Evidence* tool of GeMoMa to extract intron information from the respective RNA-seq libraries, specifying the strand specificity determined previously, and merge these from the individual libraries by the *Denoise Introns* tool of GeMoMa. This intron information may guide GeMoMa in the specific selection of splice sites during the homology-based prediction of transcript models. For predictions using GeMoMa, we specify the respective reference annotations and the files with the intron information as input of the *GeMoMa pipeline* tool. Predictions for individual species are stored for further processing in the next step.

### Merging predictions from multiple libraries

First, we merge the individual predictions of GeMoRNA used for the benchmark and we merge the individual predictions of GeMoMa based on the three reference species per target species using the *GeMoMa Annotation Filter* tool of GeMoMa (cf. Supplementary Methods).

Second, the resulting merged predictions of GeMoRNA (RNA-seq-based) and GeMoMa (homology-based) are merged using the auxiliary *Merge* tool of GeMoRNA. Here, we consider the modes “intersect”, which only reports transcript models with matching CDSs in the GeMoRNA and GeMoMa predictions, “union”, which reports the redundancy-filtered union of the GeMoRNA and GeMoMa predictions, and “intermediate”, which reports all transcript models of the “intersect” mode and additional, quality-filtered predictions from the “union” model (cf. Supplementary Methods).

### *N. benthamiana* data

We use the combination of RNA-seq-based predictions of GeMoRNA and homology-based predictions of GeMoMa to re-annotate the recently published “NbLab360” (33) and “NbKLAB” (34) strains of *Nicotiana benthamiana* (cf. Supplementary Table F). To this end, we again sampled public RNA-seq libraries of *N. benthamiana* from the European Nucleotide Archive as described in Supplementary Methods. A list of all sampled 319 libraries (ENA accessions) is provided as Supplementary Table E.

Homology-based predictions were based on the reference species *A. thaliana, S. lycopersicum* and *Nicotiana tabacum*, and based on previous *N. benthamiana* genome versions 1.0.1 and 2.6.1. For *A. thaliana* and *S. lycopersicum*, we used the genome versions listed in Supplementary Table A, while the genome versions of *N. benthamiana* and *N. tabacum* are listed in Supplementary Table F.

### Predicting transcripts in *N. benthamiana*

For predicting transcript models in the NbLab350 and NbKLAB genomes of *N. benthamiana*, we essentially follow the same procedure as previously described for the benchmark studies. Briefly, we map the RNA-seq reads to the respective genome sequence and run GeMoRNA for each of the *N. benthamiana* RNA-seq data sets separately and merge these individual predictions using the “GeMoMa Annotation Filter” tool. We also run GeMoMa for the three reference species and the two previous *N. benthamiana* genome versions and merge these as well. Finally, we merge the merged predictions of GeMoRNA and GeMoMa using the auxiliary “merge” tool of GeMoRNA. In addition to the “intersect”, “union” and “intermediate” modes, we run the “merge” tool in “annotate” mode, which results in the same set of output predictions as the union tool, but annotates transcript predictions as “high” confidence if they had been reported in the “intersect” mode, as “medium” confidence if they had been reported in the “intermediate” mode, and as “low” confidence otherwise.

## RESULTS & DISCUSSION

### Benchmark using short-read data

GeMoRNA has been benchmarked on short-read (Illumina) data for three plant species (*A. thaliana, O. sativa* and *S. lycopersicum*), three animal species (*C. elegans, D. melanogaster* and *M. musculus*), and one yeast (*S. cerevisiae*). These species have been specifically selected for their mature genome sequences and high-quality genome annotations, since these genome annotations serve as a “gold standard” in the evaluation of prediction performance.

### Evaluation on the level of transcripts

After applying Cufflinks, Scallop, StringTie and GeMoRNA to each of the RNA-seq data sets used for benchmarking (cf. Benchmark data), we evaluated their prediction performance on the level of transcripts using gffcompare (31). On the level of transcripts, gffcompare reports sensitivity and precision, which have been used to compute the respective F1-measure as shown in Figure 2.

**Fig. 2.**
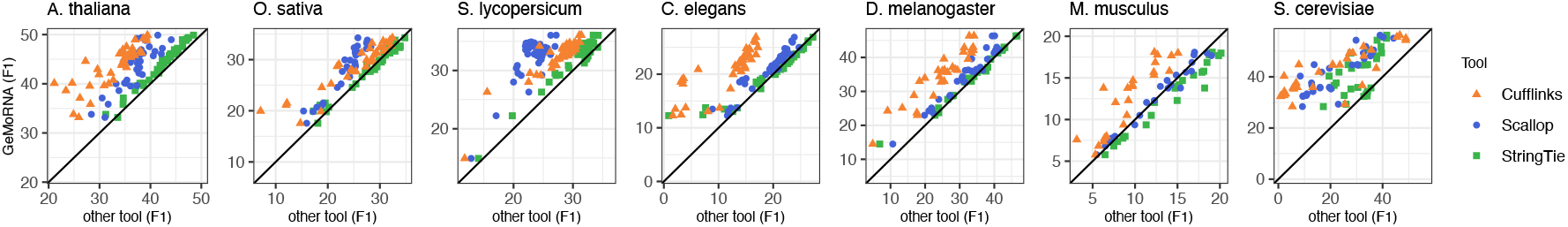
Benchmark results of GeMoRNA compared with Cufflinks, Scallop and StringTie based on the evaluation using gffcompare on the level of transcripts. Each panel displays the F1 measure based on sensitivity and precision of GeMoRNA (ordinate) compared with the remaining tools (abscissa, color coded). Points above the main diagonal indicate an improved performance of GeMoRNA, while points below the diagonal indicate an improved performance of the respective alternative tool.

In general, F1 values well below the theoretical maximum of 1.0 are to be expected, since not all genes will be expressed in a single experiment. For the majority of species, we observe that Cufflinks yields lower F1 values than the remaining tools. Cufflinks has been pioneering for the prediction of transcript models from RNA-seq data, but its development has stalled over the last years, and it has been *de facto* replaced by StringTie. The prediction performance of GeMoRNA and StringTie is widely comparable with some species-specific difference. For *A. thaliana* and *S. lycopersicum*, GeMoRNA yields slightly but consistently better F1 values than StringTie. For *O. sativa, C. elegans* and *D. melanogaster*, the prediction performance of GeMoRNA and StringTie is similar. For *M. musculus*, StringTie typically achieves higher F1 values than GeMoRNA. Scallop typically performs worse than StringTie (and GeMoRNA). For *M. musculus*, the F1 measures achieved by Scallop and GeMoRNA are widely similar. For *S. cerevisiae*, we observe the largest difference in prediction performance between GeMoRNA and all three alternative tools.

Considering sensitivity on the level of transcripts as reported by gffcompare (Supplementary Figure S5), we again observe lower sensitivity values for Cufflinks, while GeMoRNA, StringTie and Scallop achieve similar levels of sensitivity for the majority of species. For *M. musculus*, we again observe an advantage of StringTie, while for *S. cerevisiae* GeMoRNA yields substantially higher sensitivity values than any of the other tools.

Regarding precision (Supplementary Figure S6), we find a clear advantage of GeMoRNA compared with the remaining three approaches for *A. thaliana, O. sativa, S. lycopersicum* and *D. melanogaster*, while for some *M. musculus* data sets, StringTie achieves the highest precision, and for *S. cerevisiae* either StringTie or Scallop achieve improved precision values compared with GeMoRNA for a subset of data sets.

In summary, we find an advantage of GeMoRNA over StringTie, Scallop and Cufflinks especially for the plant species and yeast. For the animal species except *M. musculus*, the performance of StringTie and GeMoRNA is on the same level, while Cufflinks and Scallop perform worse. For *M. musculus*, we observe an advantage when using StringTie instead of GeMoRNA. However, due to its consistent performance, StringTie is part of many genome annotation pipelines and, hence, genome annotations may be biased towards StringTie in some cases. Improved levels of the F1 measure when using GeMoRNA are mostly due to improved precision of its predictions, while sensitivity reaches similar levels as StringTie and Scallop for the majority of species.

### Evaluation on the level of coding sequences

Since GeMoRNA has been primarily developed to complement the homology-based predictions of GeMoMa and for this reason has a focus on protein-coding genes, we further compare the performance of GeMoRNA and the three alternative approaches on the level of CDS (coding sequence) prediction. GeMoRNA makes simplistic CDS predictions (longest ORF) internally, while the other three tools predict exons but no CDSs. Hence, we apply the CDS prediction of GeMoRNA to the predictions of these tools for the purpose of benchmarking. We analyze prediction performance by the *Analyzer* tool of GeMoMa, as gffcompare does not allow for a CDS-centric evaluation. The results of this evaluation are presented in Figure 3 for the F1 measure, where we find an improvement of GeMoRNA compared with the other approaches for *A. thaliana, S. lycopersicum, D. melanogaster* and *S. cerevisiae* and, although less pronounced, for *O. sativa* and *C. elegans*. For *M. musculus*, StringTie still yields slightly better F1 values than GeMoRNA. Evaluation on the levels of sensitivity (Supplementary Figure S7) and precision (Supplementary Figure S8) supports the previous observation that the improved performance of GeMoRNA is widely due to improved levels of precision, while sensitivity values are more similar to those of the other tools, especially StringTie.

**Fig. 3.**
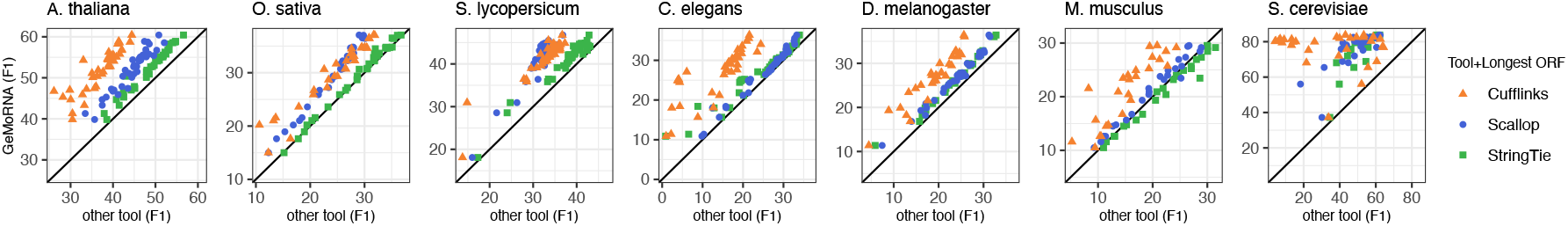
Benchmark results of GeMoRNA compared with Cufflinks, Scallop and StringTie based on the evaluation using GeMoMa Analyzer on the level of CDS. Each panel displays the F1 measure based on sensitivity and precision of GeMoRNA (ordinate) compared with the remaining tools (abscissa, color coded). Points above the main diagonal indicate an improved performance of GeMoRNA, while points below the diagonal indicate an improved performance of the respective alternative tool.

Since the GeMoRNA-internal CDS prediction based on longest ORFs within transcripts might be too simplistic and the alternative tools might profit from a more sophisticated CDS prediction, we also tested a CDS prediction using TransDecoder for these tools. For GeMoRNA, we abstain from a CDS prediction using TransDecoder since its results might contradict the GeMoRNA-internal prediction, which, however, have also been used in the split heuristics of GeMoRNA. We present the results of the evaluation based on the TransDecoder CDS prediction in Supplementary Figures S9, S10 and S11. Surprisingly, we find that the prediction performance of all tools drops considerably when using TransDecoder instead of the simplistic CDS prediction of GeMoRNA. Here, GeMoRNA yields a substantially improved prediction performance considering the F1 measure, sensitivity and precision, even for *M. musculus*. The reasons for this observation may be manifold. First, we may have used TransDecoder in an incorrect manner. However, we followed the instructions of the TransDecoder manual (cf. Supplementary Methods) and did not spot any obvious flaws. Second, the training data for TransDecoder as extracted based on predicted transcript models may not have been sufficient to accurately train the TransDecoder model. Third, the CDS annotations of the (high-quality) reference genomes may be biased towards longest ORFs, since a similar heuristic might have been used when generating these “gold standard” annotations in the first place. Based on the data available, we cannot clarify which of these options (or additional explanations) is correct. However, we consider the third explanation the most likely since, historically, the longest ORF heuristic has been applied widely.

### Assessing the influence of split heuristics

Since the improvement in prediction performance of GeMoRNA is especially pronounced for *S. cerevisiae* with its densely packed genome, one source of the improvement might be the implementation of several heuristics for splitting candidate transcripts that appear as merged on the level of read mappings. Hence, we additionally evaluate the prediction performance of a GeMoRNA variant with all split heuristics turned off. We compare GeMoRNA with and without split heuristics based on the F1 measure on the level of CDSs (Supplementary Figure S12). Indeed, we find that the improvement due to the split heuristics is especially pronounced for *S. cerevisiae*. Considerably improved F1 values can also be observed for *A. thaliana, C. elegans* and *D. melanogaster* and similar performance for the remaining three species. Hence, we may conclude that the implementation of split heuristics has an important contribution to the prediction performance of GeMoRNA, especially for dense genomes. Since these heuristics are a unique feature of GeMoRNA, it also indicates that split heuristics are partly responsible for the improvement in prediction performance compared with Cufflinks, Scallop and StringTie. However, since we also observed an improvement for *O. sativa* and *S. lycopersicum* in the previous benchmark, other aspects of the GeMoRNA algorithm (e.g., combinatorial enumeration of candidate transcripts, EM-based quantification) contribute as well, but are hard to benchmark individually, because they are deeply integrated into the complete GeMoRNA algorithm.

### Runtime and memory requirements

In addition to prediction performance, we also record the total runtime and peak memory consumption of GeMoRNA, Cufflinks, Scallop and StringTie on the benchmark data sets as measured on a compute cluster with nodes equipped with Intel Xeon E5-2680v3 processors and 128 GB of RAM. We observe (Supplementary Figure S13) that GeMoRNA requires a total runtime typically shorter than Cufflinks but longer than Scallop or StringTie. The median runtime of GeMoRNA is 14 min and 90% of the data sets are processed within 46 min. On a more recent machine (Macbook Air M3, 28 GB of RAM), absolute runtimes for these data sets are reduced to approx. 2 min (median) and 10 min (90% percentile), respectively. Regarding memory, GeMoRNA has a maximum peak memory consumption that is comparable to Scallop (55 GB vs. 52 GB), but the median memory consumption is substantially higher than for any of the other tools. One reason for the increased memory consumption is the combinatorial enumeration of candidate transcripts with associated read information. However, the median peak memory consumption of GeMoRNA is approx. 16 GB and 90% of the data sets required at most 25 GB, which makes GeMoRNA executable on current standard compute servers.

### Benchmark using long-read data

Although databases of raw sequencing data (NCBI SRA, ENA) are currently dominated by short-read, specifically Illumina, data, the amount of long-read RNA-seq data as generated by Pacific Biosciences (PacBio) or Oxford Nanopore Technologies (ONT) instruments is increasing and these will likely become the preferred technologies for the annotation of transcript models in the future. Since long RNA reads are conceptually capable of representing transcripts and transcript variants end-to-end, they are suited to improve the accuracy and completeness of genome annotations. However, the specific error profile of long-read data poses new challenges for algorithms predicting transcript models. For this reason, GeMoRNA provides a specific long-read mode that adapts some algorithmic details (cf. section 2.1). StringTie has a long-read mode as well, while Cufflinks has not been designed for long-read data and a version of Scallop (Scallop-LR (32)) specifically tailored to long-read data requires PacBio-specific header files and has not been updated for five years. Hence, we restrict the benchmark based on long-read data to the comparison of StringTie and GeMoRNA in their long-read modes. To this end, we sampled long-read data for the same seven species also considered for the short-read data and stratify these for (PacBio and ONT) instruments (cf. section Benchmarking long-read data).

On the level of transcript models evaluated using gffcompare (Supplementary Figure S14), we find an improved prediction performance of GeMoRNA for *A. thaliana, S. lycopersicum* and *C. elegans*, while for *O. sativa, D. melanogaster, M. musculus* and *S. cerevisiae*, either GeMoRNA or StringTie yields better performance for a subset of data sets, which is in some cases related to specific instrument models and/or strand specificity. This picture is widely preserved on the level of CDS using the longest-ORF heuristic for StringTie predictions (Supplementary Figure S15). Using TransDecoder for CDS prediction on the transcript models of StringTie again leads to a substantial drop in the F1 measure (Supplementary Figure S16).

### Combining RNA-seq-based and homology-based predictions

While the previous benchmarks demonstrated the accuracy of the RNA-seq-based prediction of transcripts and CDSs using GeMoRNA, the original intention for developing GeMoRNA has been to complement the homology-based predictions of GeMoMa, especially with regard to highly species-specific genes and transcripts. Hence, we evaluate the improvements of annotation completeness and accuracy combining GeMoMa and GeMoRNA predictions in the following. To this end, we additionally perform homology-based transcript prediction using GeMoMa for the seven species considered in the benchmark studies (cf. Methods, Reference species listed in Supplementary Table D). We first merge all RNA-seq-based GeMoRNA predictions based on the data sets used for the benchmark, and we merge all homology-based GeMoMa predictions across the reference species considered. Afterwards, the individually merged GeMoRNA and GeMoMa predictions are further combined by determining the intersection and union of the sets of predictions of both tools per species. We might expect that the intersection of independent sources of evidence (homology and RNA-seq) yields high precision, while the union of both sources would achieve high sensitivity. We additionally consider an “intermediate” variant that supplements the intersection set with further high confidence predictions of the individual tools (cf. Methods). First, we consider sensitivity on the level of CDSs of the individual prediction sets of GeMoRNA and GeMoMa, and the different combinations of these. In Supplementary Figure S17, we indeed observe that the union of the predictions of both tools yields the best sensitivity. However, the improvement over the individual predictions of GeMoRNA and GeMoMa is lower than we expected, and the intersection of both sets results in only mildly reduced sensitivity. This indicates that the overlap between the RNA-seq-based predictions of GeMoRNA and the homology-based predictions of GeMoMa is substantial, and may provide a set of high-confidence transcript models. As expected, we find the largest precision values for the intersection of GeMoRNA and GeMoMa predictions, while precision drops substantially for the union of both (Figure 4). The intermediate variant of the combination yields a considerably improved precision compared with the individual predictions except for *S. cerevisiae*, for which we observe precision values above 75% in all cases. Notably, we observe precision values above 85% for the intersection for four of the species (*A. thaliana, C. elegans, M. musculus, S. cerevisiae*), while precision is substantially lower for the remaining species. Since the intersection of GeMoRNA and GeMoMa predictions provides evidence of the respective transcript models from two independent sources (RNA-seq and homology), we assume the predictions in the intersection to be likely correct. Hence, this finding may indicate that the current reference annotation of those species (*O. sativa, S. lycopersicum, D. melanogaster*) is less complete than for the other species. Turning to the F1 measure combining sensitivity and precision (Supplementary Figure S18), we find that – as intended – the intermediate variant for combing GeMoRNA and GeMoMa predictions yields the best F1 values for four of the species (*A. thaliana, C. elegans, D. melanogaster, M. musculus*). For the remaining species, F1 values are comparable to the intersection case, that, however, achieved substantially lower sensitivity, which might be relevant for practical applications.

**Fig. 4.**
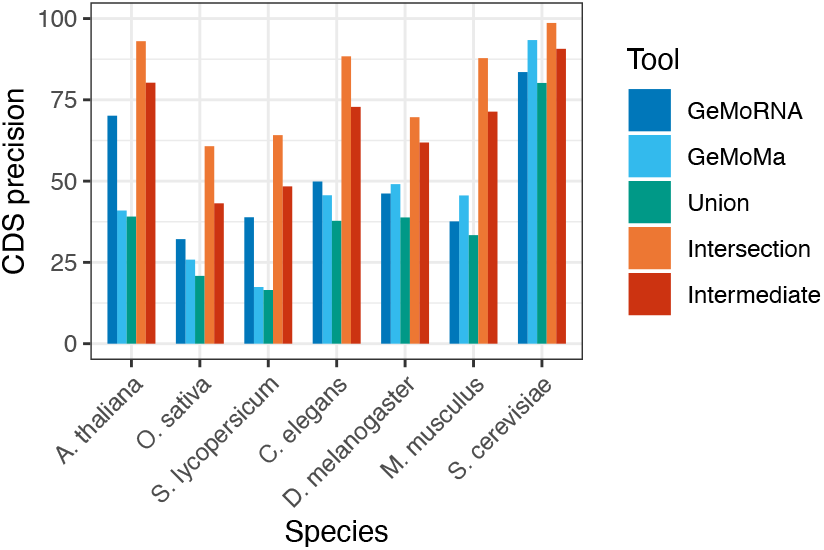
Precision on the level of CDS for GeMoRNA (RNA-seq-based) and GeMoMa (homology-based) predictions, and for combinations of the prediction sets.

### Applying GeMoRNA to *N. benthamiana* LAB genomes

*Nicotiana benthamiana* is used as a model plant in many experimental labs worldwide, especially since it is conveniently infiltrated with *Agrobacterium tumefaciens* for transient expression of genes. Recently, two genomes and corresponding annotation of “lab” strains have been published (33, 34). On first sight, the transcript annotations of these genomes appear to be rather incomplete with mostly one or only few transcript variants per gene. In addition, both transcript annotations use transcript IDs that are unrelated to the previously used *N. benthamiana* v1.0.1 and v2.6.1 genomes, which complicates the comparisons with previous publications. Hence, we set out to apply the combination of RNA-seq-based GeMoRNA predictions and homology-based GeMoMa predictions to both lab genomes. For GeMoRNA, we compile a collection of 319 *N. benthamiana* RNA-seq data sets from the European Nucleotide Archive (cf. Methods, Supplementary Table E). For GeMoMa, we consider *A. thaliana, S. lycopersicum* and *N. tabacum* as reference species and additionally include the previous v.1.0.1 and v.2.6.1 genome versions to establish the link between this new annotation and the previous ones. We merge the GeMoRNA and GeMoMa predictions as described for the benchmark studies, and assign confidence levels “high”, “medium” and “low” as described in Methods. In the merge, transcript IDs of the v1.0.1 and v2.6.1 are available either in the ID field or as entries of the “alternative” tag of the GFF format if these could be transferred to the respective lab genome.

In Figure 5A, we show an upset plot (inclusive intersection size) of the NbLab360 v103 annotation and the annotation generated using GeMoRNA and GeMoMa on different levels of confidence. The NbLab360 annotation contains 45,796 (protein coding) transcripts, while the high-confidence intersection of GeMoRNA and GeMoMa contains 35,684 transcripts, 21,193 of which are shared (i.e., identical on the level of CDSs) between both annotations. Also considering medium-confidence GeMoRNA/GeMoMa predictions, the number of transcripts increases to 65,892 belonging to 43,307 genes, which corresponds to 1.78 transcript per gene on average. A number of 26,874 of these transcripts is shared with the NbLab360 annotation. Also including low-confidence transcripts, the number of transcripts increases dramatically to 231,219 transcripts. Hence, we recommend to consider low-confidence transcripts only when aiming for maximum sensitivity.

**Fig. 5.**
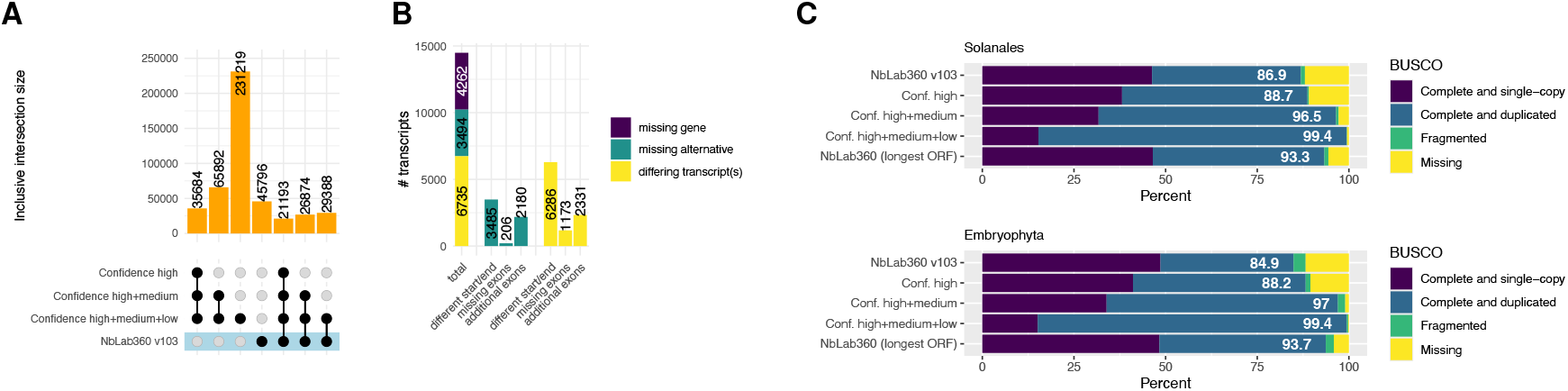
Comparison of the original NbLab360 annotation to the merged prediction of GeMoRNA and GeMoMa considering transcripts of different levels of confidence. (A) Intersection sizes (inclusive, i.e., size of intersection shown irrespective of the occurrence in other sets/intersections) of the CDS contained in each of the annotation files. (B) Breakdown of transcripts that are present in the “Confidence high” set of GeMoRNA/GeMoMa but are missing from the NbLab360 annotation. (C) BUSCO completeness of the annotation files (extracted proteins) for the solanales and embryophyta data sets.

We further scrutinize the 14,491 transcripts that are present in the high-confidence GeMoRNA/GeMoMa set but not in the NbLab360 annotation (Figure 5B). We find that 4,262 of these transcript do not overlap an existing NbLab360 annotation and may, hence, be considered transcripts of missing genes. If a GeMoRNA/GeMoMa transcript does not match any NbLab360 transcript, but another GeMoRNA/GeMoMa transcript of the same gene perfectly matches an NbLab360 transcript, these GeMoRNA/GeMoMa transcripts may be considered missing alternative transcripts. We find 3,494 of such missing alternatives, which more frequently contain additional exons (2,180 cases) than they lack exons (206 cases) present in the matching variant. GeMoRNA/GeMoMa transcripts that overlap but do not perfectly match any NbLab360 transcript may differ in multiple aspects, where alternative transcript start or end positions are the most frequent (6,286 out of 6,735 differing transcripts).

For the transcripts from the “missing gene” category, we aim at obtaining an overview of the functions that are encoded by these genes. To this end, we extract the protein sequences of all high-confidence transcripts and use PANNZER2 (35) to assign functional annotation and GO terms. Among the transcripts from the “missing gene” category, the terms (biological product) “regulation of DNA-templated transcription” (161 transcripts), “defense response” (116), “translation” (103), “protein ubiquitination” (82), “chromatin remodeling” (80), “response to auxin” (67) and “methylation” (55) are the most frequent. Terms with significant enrichment (Fisher’s exact test, adjusted p-value *<* 0.05) are “killing of cells of another organism” (9 transcripts; log-odds ratio 3.6; *p* = 1.3 10^*−*2^), “cell population proliferation” (11; 3.5; *p* = 1.4 10^*−*3^), “hydrotropism” (10; 3.0; *p* = 4.3 10^*−*2^), “photosynthesis, light harvesting” (17; 2.2; *p* = 9.0 10^*−*3^) and “response to auxin” (67; 1.9; *p* = 1.8 10^*−*12^). Together, this indicates that the genes that are only present in our reannotation of the NbLab genome fulfill important, central functions for *N. benthamiana* plants.

Since no established “gold standard” for *N. benthamiana* transcripts has been established, the quality of transcript models cannot be evaluated in the same manner as for the benchmark species. Hence, we resort to an assessment of completeness of the different annotation variants using BUSCO (36, 37) on the level of protein sequences. Here, we consider only CDS annotations with proper start and stop codons and without premature stops. In Figure 5C, we present the BUSCO completeness of the original NbLab360 v103 annotation and GeMoRNA/GeMoMa-derived annotations on different levels of confidence using the “Solanales” and “Embroyphyta” BUSCO sets. We observe that even the high-confidence predictions of GeMoRNA/GeMoMa achieve a better BUSCO completeness (single-copy and duplicated) than the original NbLab360 annotation, while completeness further increases when also considering medium and low-confidence transcripts. In previous benchmark studies, we noted that the longest-ORF heuristic of GeMoRNA typically results in better prediction performance than using TransDecoder. Hence, we additionally apply this heuristic to the NbLab360 v103 annotation, which may also repair previously missing start and stop codons. Indeed, we find that this increases BUSCO completeness, although completeness is still lower than for the combination of high-confidence and medium-confidence GeMoRNA/GeMoMA predictions.

We also observe that the number of duplicated BUSCOs for GeMoRNA/GeMoMa predictions is substantially larger than for the original NbLab360 v103 annotation, which contains one transcript variant per gene. Hence, we aggregate the BUSCO results on the level of genes using the “BUSCORe-computer’ module of GeMoMa. We find (Supplementary Figure S20 and S21) that this indeed decreases the number of duplicated BUSCOs for the GeMoRNA/GeMoMa predictions to similar levels as for the NbLab360 v103 annotation, except for the low-confidence set.

We apply the same procedure to the NbKLAB genome and corresponding annotation (Supplementary Figure S19) with similar results, but a considerably larger number of missing genes (6,700) and slightly lower BUSCO completeness of the NbKLAB annotation. One notable difference is that applying the longest-ORF heuristic to the NbKLAB annotation does not result in different BUSCO completeness, likely because a similar heuristic has already been applied for CDS prediction initially and the number of missing start/stop codons is already substantially lower for the original NbKLAB annotation than for NbLab360.

## Conclusion

In benchmark studies, we found that GeMoRNA yields an improved prediction performance compared with the previous approaches Cufflinks, Scallop and StringTie for six out of seven species considered. This improvement is especially pronounced for the two plant species *A. thaliana* and *S. lycopersicum* and for yeast (*S. cerevisiae*), which have rather dense genomes. We further demonstrated how the combination of RNA-seq-based predictions of GeMoRNA with homology-based predictions using GeMoMa may be used to balance sensitivity and precision in the annotation of transcript models. Applying the combination of GeMoRNA and GeMoMa to recently sequenced genomes of two *N. benthamiana* lab strains, we illustrated the utility of this strategy to yield high-quality annotations of transcript models and coding sequences for newly sequenced genomes. We provide the GeMoRNA tool and the generated annotations of *N. benthamiana* lab strains to the scientific community.

## Data availability

The source code of GeMoRNA is available from GitHub at https://github.com/Jstacs/Jstacs/tree/master/projects/ gemorna. A binary version of GeMoRNA is available from https://www.jstacs.de/index.php/GeMoRNA. The annotation files for the *N. benthamiana* labs strains are available from zenodo at https://doi.org/10.5281/zenodo.14901380.

## ACKNOWLEDGEMENTS

This preprint is formatted using a LATEX class by Ricardo Henriques. We thank Lilya Kopertekh for valuable discussions.

