## Supplementary Methods, Tables A, D, F, Figures S1-S21 for "Improved reconstruction of transcripts and coding sequences from RNA-seq data"

### SUPPLEMENTARY MATERIAL

#### Supplementary Methods – transcript-specific and read-specific weights

*A priori* transcript weights are defined as

$$w_t = \frac{\ell_t}{\sum_{\tilde{t}\in T} \ell_{\tilde{t}}},$$

where $\ell_t$ denotes the length of the CDS (longest ORF) of candidate transcript $t$, and shall favor transcript models with long, intact CDS predictions.
Let $n_i$ denote the number of reads supporting intron $i$ of the splicing graph, and let $m_e$ denote the number of reads supporting exon $e$. Let further denote $I(r)$ the set of introns that are supported by read $r$ and $E(r)$ denote the set of exons that are supported by read $r$. Let $\bar{n}_i$ denote the maximum number of supporting reads of any exon overlapping intron $i$ (i.e., exons of other candidate transcripts, which are alternatives to that intron), or $\bar{n}_i = 1$ if no such exons exist. In analogy, let $\bar{m}_e$ denote the maximum number of supporting reads of any intron overlapping exon $e$.
We define

$$N_r = \prod_{i\in I(r)} \frac{2n_i}{\bar{n}_i + n_i}$$

$$M_r = \prod_{e\in E(r)} \frac{2m_e}{\bar{m}_e + m_e}$$

and based on these quantities set the weight of read $r$ to

$$v_r = (N_r \cdot M_r)^{\frac{1}{|I(r)|+|E(r)|}} \cdot s^{|I(r)|},$$

where $s = 5$. The first two factors shall put greater weight on reads that help to distinguish between different transcript variants (alternative exons and introns) while the latter puts greater weight on reads spanning (multiple) introns, in general.
For long-read data, the weighting of reads in the quantification process is adapted to

$$v_r = (N_r \cdot M_r)^{\frac{1}{|I(r)|+|E(r)|}} \cdot L_{|I(R)|},$$

where $L_0 = 1$, $L_1 = s$, $L_n = L_{n-1} \cdot \left(1 + \left(\frac{2}{3}\right)^{n-1}(s-1)\right)$, which results in an exponentially decreasing additional weight contribution from additional introns.

#### Supplementary Methods – TransDecoder

When applying TransDecoder, we followed the instructions of the TransDecoder wiki (https://github.com/TransDecoder/TransDecoder/wiki) by performing the following steps:

- extract transcript FastA files using
```
gtf_genome_to_cdna_fasta.pl <gtf> \
  <genome> > transcripts.fasta
```
- convert the GTF/GFF to the internal "alignment" GFF
```
gtf_to_alignment_gff3.pl <gtf> \
  > transcripts.gff3
```
- train and apply the TransDecoder models
```
TransDecoder.LongOrfs \
  -t transcripts.fasta -S
TransDecoder.Predict \
  -t transcripts.fasta
```
- convert the results back to a genomic GFF
```
cdna_alignment_orf_to_genome_orf.pl \
  transcripts.fasta.transdecoder.gff3 \
  transcripts.gff3
  transcripts.fasta \
  > transdecoder.genome.gff3
```

##### Supplementary Methods – GeMoMa Annotation Filter

For the GeMoRNA predictions, we specify the filter criteria of the *GeMoMa Annotation Filter* tool as `start='M' and stop='*' and sumWeight>1`, which filters the merged transcript models for those that have a proper start and stop codon and have evidence from at least two RNA-seq libraries. In addition, we specify the alternative transcript filter as `sumWeight>2 and aa>=30 and ce>1`, which filters additional transcript models of a common gene for those that have evidence from at least three RNA-seq libraries, encode a protein with at least 30 amino acids and have at least two coding exons.

For the GeMoMa predictions, we leave the filter criteria at their default values and specify the alternative transcript filter as `(tie==1 or sumWeight>1) and aa>=30 and ce>1`, which filters additional transcript models of a common gene for those that have support from RNA-seq data for all their splice sites or have been predicted based on at least two reference species, encode a protein with at least 30 amino acids and have at least two coding exons.

For quality filtering, we perform additional runs of *GeMoMa Annotation Filter* as described previously, but additional, stringent filter criteria, `sumWeight>4 and tie==1.0` in case of GeMoRNA and `score/aa>=1.0 and tie==1.0` in case of GeMoMa predictions.

##### Supplementary Methods – sampling of *N. benthamiana* data sets

We queried all data sets for *N. benthamiana* with `library_layout` "paired", `library_source` "transcriptomic" and `library_strategy` "RNA-Seq", which yielded a list of $1{,}413$ data sets. From these, we selected all data sets that have been used in the original "NbLab360" annotation (study accession PRJNA881799). In addition, we considered a sample of data sets for each study accession present in ENA. If multiple sequencing runs existed for a specific biosample, we considered the run with the largest number of reads. From these, we sampled a subset of runs for each study accession. I less than 5 runs existed for a specific accession, all runs were considered. For study accessions with a larger number of runs, 5 of these were sampled at random. This procedure resulted in a collection of 319 *N. benthamiana* RNA-seq data sets that were analyzed further.

**Supplementary Table A.** List of sources of genome sequence and annotation for species considered in benchmark studies.

| | |
|---|---|
| Species | *A. thaliana* |
| Source | Ensembl |
| Genome | https://ftp.ensemblgenomes.ebi.ac.uk/pub/plants/release-56/fasta/arabidopsis_thaliana/dna/Arabidopsis_thaliana.TAIR10.dna.toplevel.fa.gz |
| Annotation | https://ftp.ensemblgenomes.ebi.ac.uk/pub/plants/release-56/gff3/arabidopsis_thaliana/Arabidopsis_thaliana.TAIR10.56.gff3.gz |
| Species | *O. sativa* |
| Source | Ensembl |
| Genome | https://ftp.ensemblgenomes.ebi.ac.uk/pub/plants/release-56/fasta/oryza_sativa/dna/Oryza_sativa.IRGSP-1.0.dna.toplevel.fa.gz |
| Annotation | https://ftp.ensemblgenomes.ebi.ac.uk/pub/plants/release-56/gff3/oryza_sativa/Oryza_sativa.IRGSP-1.0.56.gff3.gz |
| Species | *S. lycopersicum* |
| Source | solgenomics.net |
| Genome | https://solgenomics.net/ftp/tomato_genome/Heinz1706/assembly/build_4.00/S_lycopersicum_chromosomes.4.00.fa.gz |
| Annotation | https://solgenomics.net/ftp/tomato_genome/Heinz1706/annotation/ITAG4.1_release/ITAG4.1_gene_models.gff |
| Species | *C. elegans* |
| Source | wormbase.org |
| Genome | https://downloads.wormbase.org/species/c_elegans/sequence/genomic/c_elegans.PRJEB28388.WS287.genomic.fa.gz |
| Annotation | https://downloads.wormbase.org/species/c_elegans/gff/c_elegans.PRJEB28388.WS287.annotations.gff3.gz |
| Species | *D. melanogaster* |
| Source | flybase.net |
| Genome | http://ftp.flybase.net/genomes/Drosophila_melanogaster/dmel_r6.50_FB2023_01/fasta/dmel-all-chromosome-r6.50.fasta.gz |
| Annotation | http://ftp.flybase.net/genomes/Drosophila_melanogaster/dmel_r6.50_FB2023_01/gff/dmel-all-r6.50.gff.gz |
| Species | *M. musculus* |
| Source | NCBI |
| Genome | https://ftp.ncbi.nlm.nih.gov/genomes/all/GCF/000/001/635/GCF_000001635.27_GRCm39/GCF_000001635.27_GRCm39_genomic.fna.gz |
| Annotation | https://ftp.ncbi.nlm.nih.gov/genomes/all/GCF/000/001/635/GCF_000001635.27_GRCm39/GCF_000001635.27_GRCm39_genomic.gff.gz |
| Species | *S. cerevisiae* |
| Source | NCBI |
| Genome | https://ftp.ncbi.nlm.nih.gov/genomes/all/GCF/000/146/045/GCF_000146045.2_R64/GCF_000146045.2_R64_genomic.fna.gz |
| Annotation | https://ftp.ncbi.nlm.nih.gov/genomes/all/GCF/000/146/045/GCF_000146045.2_R64/GCF_000146045.2_R64_genomic.gff.gz |

**Supplementary Table B.** List of data sets used for benchmarking.

[Available as separate file `Illumina_datasets.xls`]

**Supplementary Table C.** List of long-read data sets used for benchmarking.

[Available as separate file `Long_read_datasets.xlsx`]

**Supplementary Table D.** List of reference species and genome versions used for homology-based prediction with GeMoMa.

| | |
|---|---|
| ***A. thaliana*** | |
| Species | *Brassica rapa* |
| Source | BnPIR |
| Genome | http://cbi.hzau.edu.cn/rape/download_ext/zs11.genome.fa |
| Annotation | http://cbi.hzau.edu.cn/rape/download_ext/zs11.v0.gff3 |
| Species | *Capsella rubella* |
| Source | Phytozome |
| Genome | `Crubella_474_v1.fa` |
| Annotation | `Crubella_474_v1.1.gene_exons.gff3` |
| Species | *Oryza sativa* |
| Source | Ensembl |
| Genome | https://ftp.ensemblgenomes.ebi.ac.uk/pub/plants/release-56/fasta/oryza_sativa/dna/Oryza_sativa.IRGSP-1.0.dna.toplevel.fa.gz |
| Annotation | https://ftp.ensemblgenomes.ebi.ac.uk/pub/plants/release-56/gff3/oryza_sativa/Oryza_sativa.IRGSP-1.0.56.gff3.gz |
| ***O. sativa*** | |
| Species | *Arabidopsis thaliana* |
| Source | Ensembl |
| Genome | https://ftp.ensemblgenomes.ebi.ac.uk/pub/plants/release-56/fasta/arabidopsis_thaliana/dna/Arabidopsis_thaliana.TAIR10.dna.toplevel.fa.gz |
| Annotation | https://ftp.ensemblgenomes.ebi.ac.uk/pub/plants/release-56/gff3/arabidopsis_thaliana/Arabidopsis_thaliana.TAIR10.56.gff3.gz |
| Species | *Brachypodium distachyon* |
| Source | Phytozome |
| Genome | `Bdistachyon_314_v3.0.fa.gz` |
| Annotation | `Bdistachyon_314_v3.1.gene.gff3.gz` |
| Species | *Zizania latifolia* |
| Source | CNCB-NGDC |
| Genome | https://download.cncb.ac.cn/gwh/Plants/Zizania_latifolia_Zlat_genome_v1_GWHBFHI00000000/GWHBFHI00000000.genome.fasta.gz |
| Annotation | https://download.cncb.ac.cn/gwh/Plants/Zizania_latifolia_Zlat_genome_v1_GWHBFHI00000000/GWHBFHI00000000.gff.gz |
| ***S. lycopersicum*** | |
| Species | *Arabidopsis thaliana* |
| Source | Ensembl |
| Genome | https://ftp.ensemblgenomes.ebi.ac.uk/pub/plants/release-56/fasta/arabidopsis_thaliana/dna/Arabidopsis_thaliana.TAIR10.dna.toplevel.fa.gz |
| Annotation | https://ftp.ensemblgenomes.ebi.ac.uk/pub/plants/release-56/gff3/arabidopsis_thaliana/Arabidopsis_thaliana.TAIR10.56.gff3.gz |
| Species | *Nicotiana benthamiana* |
| Source | solgenomics.net |
| Genome | https://solgenomics.net/ftp/genomes/Nicotiana_benthamianaV261/Nbenthamiana_Assembly/Niben261_genome.fasta.gz |
| Annotation | https://solgenomics.net/ftp/genomes/Nicotiana_benthamianaV261/Nbenthamiana_Annotation/Niben261_genome.annotation.gene_models.gff.gz |
| Species | *Solanum tuberosum* |
| Source | solgenomics.net |
| Genome | https://solgenomics.net/organism/Solanum_tuberosum/genome: `PGSC_DM_v3_superscaffolds.fasta` |
| Annotation | https://solgenomics.net/organism/Solanum_tuberosum/genome: `PGSC_DM_v3.4_gene.gff` |
| ***C. elegans*** | |
| Species | *Caenorhabditis japonica* |
| Source | wormbase.org |
| Genome | https://downloads.wormbase.org/species/c_japonica/sequence/genomic/c_japonica.PRJNA12591.WS288.genomic.fa.gz |

| | |
|---|---|
| Annotation | https://downloads.wormbase.org/species/c_japonica/gff/c_japonica.PRJNA12591.WS288.annotations.gff3.gz |
| Species | *Caenorhabditis briggsae* |
| Source | wormbase.org |
| Genome | https://downloads.wormbase.org/species/c_briggsae/sequence/genomic/c_briggsae.PRJNA10731.WS288.genomic.fa.gz |
| Annotation | https://downloads.wormbase.org/species/c_briggsae/gff/c_briggsae.PRJNA10731.WS288.annotations.gff3.gz |
| Species | *Caenorhabditis brenneri* |
| Source | wormbase.org |
| Genome | https://downloads.wormbase.org/species/c_brenneri/sequence/genomic/c_brenneri.PRJNA20035.WS288.genomic.fa.gz |
| Annotation | https://downloads.wormbase.org/species/c_brenneri/gff/c_brenneri.PRJNA20035.WS288.annotations.gff3.gz |
| ***D. melanogaster*** | |
| Species | *Drosophila simulans* |
| Source | flybase.net |
| Genome | http://ftp.flybase.net/genomes/Drosophila_simulans/dsim_r2.02_FB2017_04/fasta/dsim-all-chromosome-r2.02.fasta.gz |
| Annotation | http://ftp.flybase.net/genomes/Drosophila_simulans/dsim_r2.02_FB2017_04/gff/dsim-all-r2.02.gff.gz |
| Species | *Drosophila grimshawi* |
| Source | flybase.net |
| Genome | http://ftp.flybase.net/genomes/Drosophila_grimshawi/dgri_r1.3_FB2016_05/fasta/dgri-all-chromosome-r1.3.fasta.gz |
| Annotation | http://ftp.flybase.net/genomes/Drosophila_grimshawi/dgri_r1.3_FB2016_05/gff/dgri-all-r1.3.gff.gz |
| Species | *Ceratitis capitata* |
| Source | NCBI |
| Genome | https://ftp.ncbi.nlm.nih.gov/genomes/all/GCF/000/347/755/GCF_000347755.3_Ccap_2.1/GCF_000347755.3_Ccap_2.1_genomic.fna.gz |
| Annotation | https://ftp.ncbi.nlm.nih.gov/genomes/all/GCF/000/347/755/GCF_000347755.3_Ccap_2.1/GCF_000347755.3_Ccap_2.1_genomic.gff.gz |
| ***M. musculus*** | |
| Species | *Rattus norvegicus* |
| Source | NCBI |
| Genome | https://ftp.ncbi.nlm.nih.gov/genomes/all/GCF/015/227/675/GCF_015227675.2_mRatBN7.2/GCF_015227675.2_mRatBN7.2_genomic.fna.gz |
| Annotation | https://ftp.ncbi.nlm.nih.gov/genomes/all/GCF/015/227/675/GCF_015227675.2_mRatBN7.2/GCF_015227675.2_mRatBN7.2_genomic.gff.gz |
| Species | *Mesocricetus auratus* |
| Source | NCBI |
| Genome | https://ftp.ncbi.nlm.nih.gov/genomes/all/GCF/017/639/785/GCF_017639785.1_BCM_Maur_2.0/GCF_017639785.1_BCM_Maur_2.0_genomic.fna.gz |
| Annotation | https://ftp.ncbi.nlm.nih.gov/genomes/all/GCF/017/639/785/GCF_017639785.1_BCM_Maur_2.0/GCF_017639785.1_BCM_Maur_2.0_genomic.gff.gz |
| Species | *Homo sapiens* |
| Source | NCBI |
| Genome | https://ftp.ncbi.nlm.nih.gov/genomes/all/GCF/000/001/405/GCF_000001405.40_GRCh38.p14/GCF_000001405.40_GRCh38.p14_genomic.fna.gz |
| Annotation | https://ftp.ncbi.nlm.nih.gov/genomes/all/GCF/000/001/405/GCF_000001405.40_GRCh38.p14/GCF_000001405.40_GRCh38.p14_genomic.gff.gz |
| ***S. cerevisiae*** | |
| Species | *Saccharomyces paradoxus* |
| Source | NCBI |
| Genome | https://ftp.ncbi.nlm.nih.gov/genomes/all/GCF/002/079/055/GCF_002079055.1_ASM207905v1/GCF_002079055.1_ASM207905v1_genomic.fna.gz |

| | |
|---|---|
| Annotation | https://ftp.ncbi.nlm.nih.gov/genomes/all/GCF/002/079/055/GCF_002079055.1_ASM207905v1/GCF_002079055.1_ASM207905v1_genomic.gff.gz |
| Species | *Schizosaccharomyces pombe* |
| Source | NCBI |
| Genome | https://ftp.ncbi.nlm.nih.gov/genomes/all/GCF/000/002/945/GCF_000002945.1_ASM294v2/GCF_000002945.1_ASM294v2_genomic.fna.gz |
| Annotation | https://ftp.ncbi.nlm.nih.gov/genomes/all/GCF/000/002/945/GCF_000002945.1_ASM294v2/GCF_000002945.1_ASM294v2_genomic.gff.gz |
| Species | *Kluyveromyces lactis* |
| Source | NCBI |
| Genome | https://ftp.ncbi.nlm.nih.gov/genomes/all/GCF/000/002/515/GCF_000002515.2_ASM251v1/GCF_000002515.2_ASM251v1_genomic.fna.gz |
| Annotation | https://ftp.ncbi.nlm.nih.gov/genomes/all/GCF/000/002/515/GCF_000002515.2_ASM251v1/GCF_000002515.2_ASM251v1_genomic.gff.gz |

**Supplementary Table E.** List of data sets used for predicting transcript models in *N. benthamiana*.

[Available as separate file `N_benthamiana_datasets.xlsx`]

**Supplementary Table F.** List of *N. benthamiana* genome version and *N. tabacum* genome used for homology-based prediction of transcripts.

| | |
|---|---|
| Species | *N. benthamiana* (NbLab360) |
| Source | nbenth.com |
| Genome | https://bioweb01.qut.edu.au/N.bethamiana/NbLab360.genome.fasta.gz |
| Annotation | https://bioweb01.qut.edu.au/N.bethamiana/NbLab360.v103.gff3.gz |
| Species | *N. benthamiana* (NbKLAB) |
| Source | figshare.com |
| Genome | https://figshare.com/ndownloader/files/45147532 |
| Annotation | https://figshare.com/ndownloader/files/45147526 |
| Species | *N. benthamiana* (v1.0.1) |
| Source | solgenomics.net |
| Genome | https://solgenomics.net/ftp/genomes/Nicotiana_benthamiana/assemblies/Niben.genome.v1.0.1.scaffolds.nrcontigs.fasta.gz |
| Annotation | https://solgenomics.net/ftp/genomes/Nicotiana_benthamiana/annotation/Niben101/Niben101_annotation.gene_models.gff |
| Species | *N. benthamiana* (v2.6.1) |
| Source | solgenomics.net |
| Genome | https://solgenomics.net/ftp/genomes/Nicotiana_benthamianaV261/Nbenthamiana_Assembly/Niben261_genome.fasta.gz |
| Annotation | https://solgenomics.net/ftp/genomes/Nicotiana_benthamianaV261/Nbenthamiana_Annotation/Niben261_genome.annotation.gene_models.gff.gz |
| Species | *N. tabacum* |
| Source | solgenomics.net |
| Genome | https://solgenomics.net/ftp/genomes/Nicotiana_tabacum/edwards_et_al_2017/assembly/Nitab-v4.5_genome_Scf_Edwards2017.fasta.gz |
| Annotation | https://solgenomics.net/ftp/genomes/Nicotiana_tabacum/edwards_et_al_2017/annotation/Nitab-v4.5_gene_models_Scf_Edwards2017.gff |

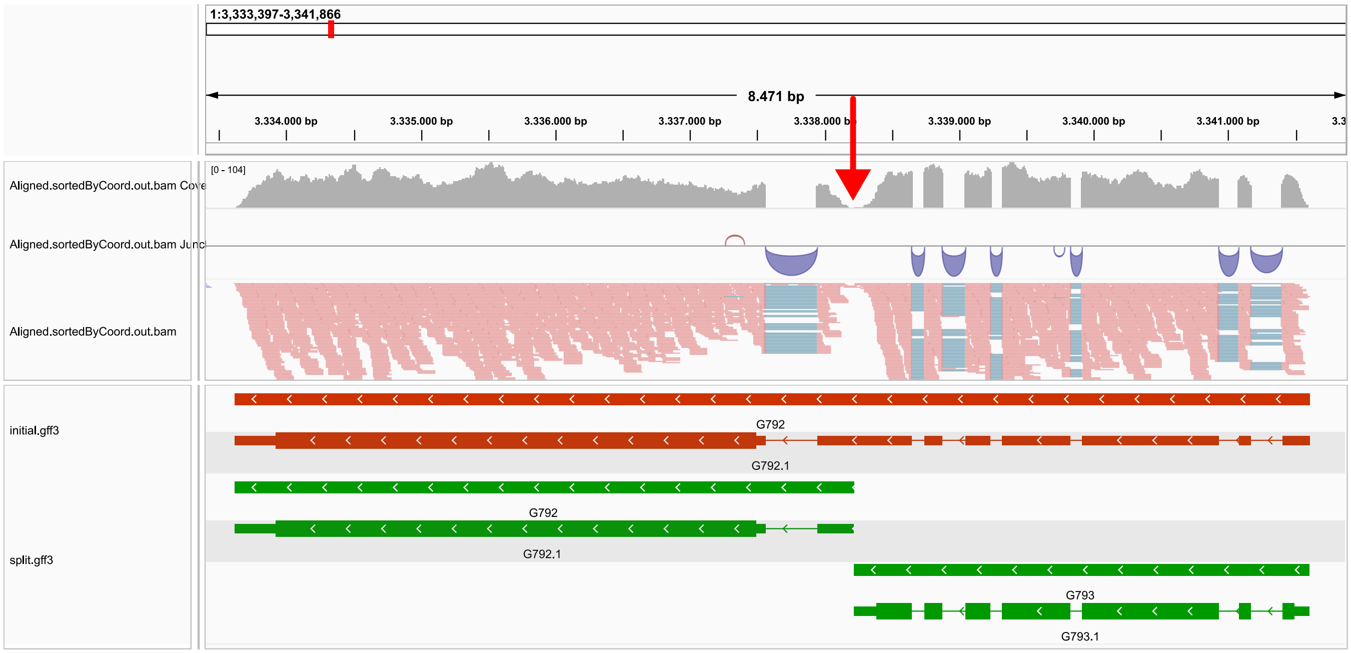


**Supplementary Figure S1.** Illustration of the split heuristic based on local dips in the coverage profile. Considering the mapped reads, we observe contiguous coverage in this region, which results in one initial candidate transcript (red). Within the coverage profile, we find one pronounced dip (red arrow), which is considered as a possible split position. Since a split at this position does not affect the original CDS prediction (thicker bar in red transcript) and both resulting split transcript models (green) obtain a CDS prediction, a split is performed.

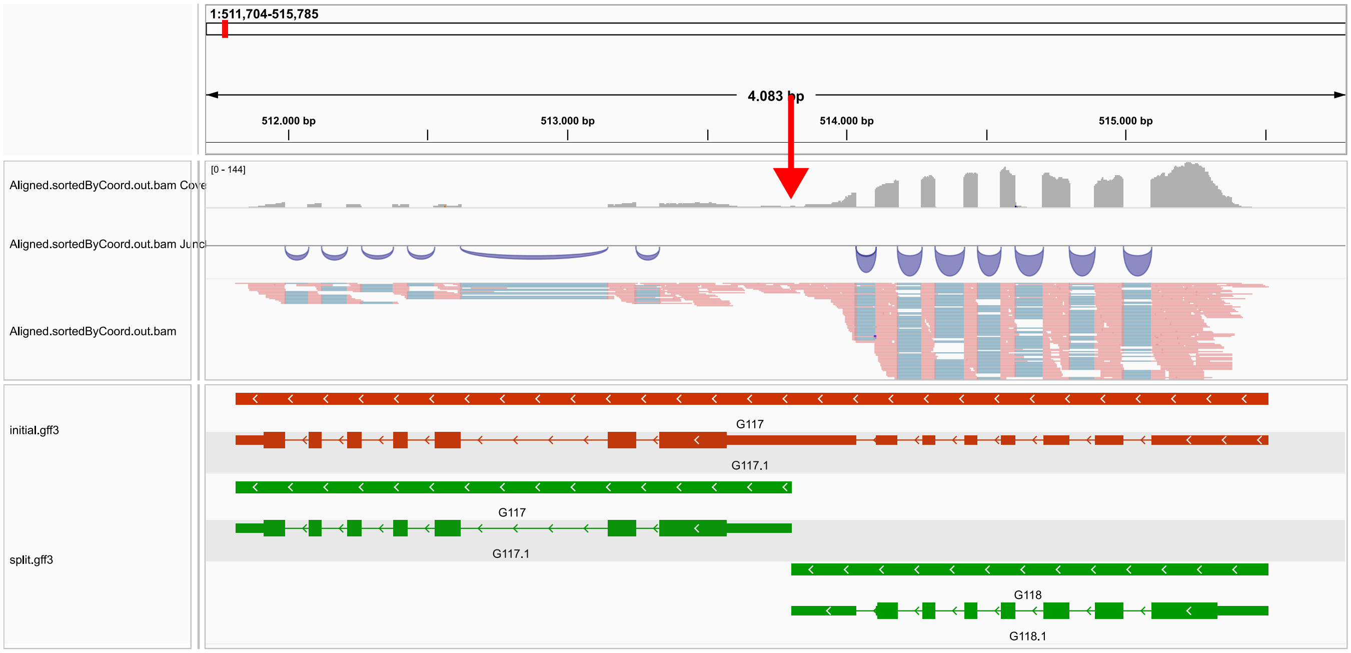


**Supplementary Figure S2.** Illustration of the split heuristic based on a pronounced coverage imbalance in the coverage profile. Considering the mapped reads, we observe contiguous coverage in this region, which results in one initial candidate transcript (red). In the coverage profile, we find an imbalance with low coverage to the left and higher coverage to the right of the red arrow, which is considered as a possible split position. Since a split at this position does not affect the original CDS prediction (thicker bar in red transcript) and both resulting split transcript models (green) obtain a CDS prediction, a split is performed.

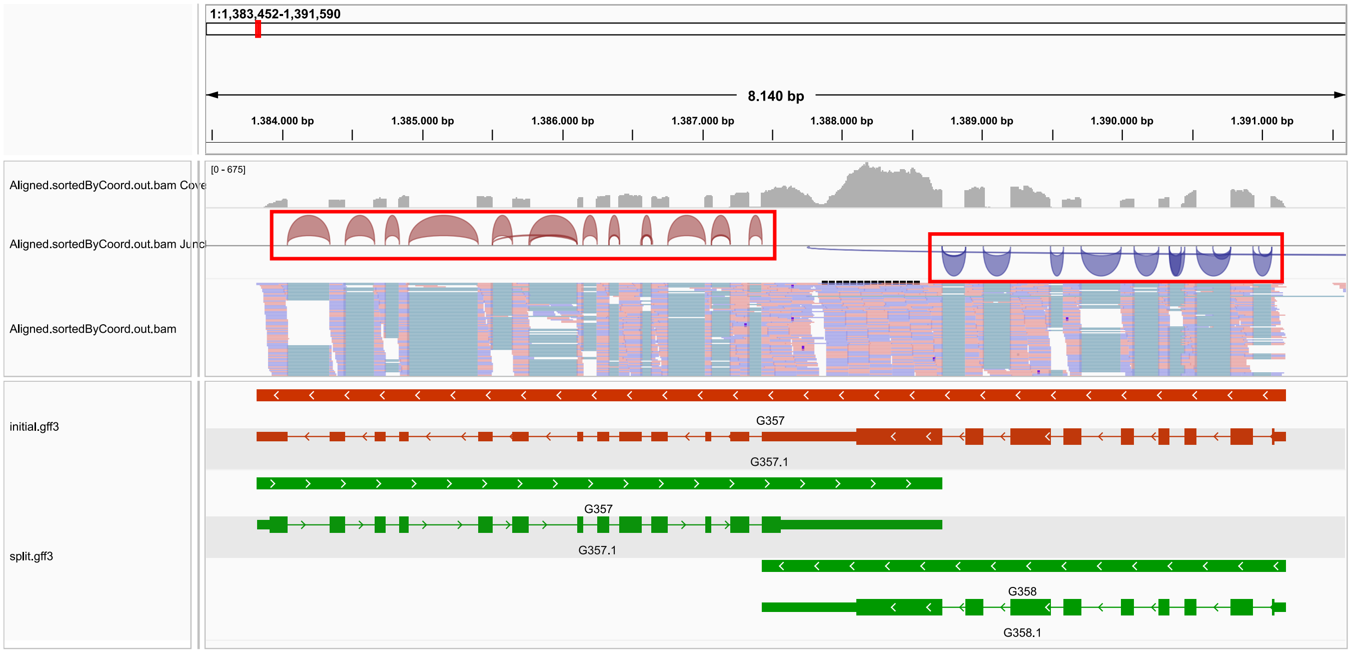


**Supplementary Figure S3.** Illustration of the split heuristic based on the orientation of splice sites. As indicated by the colouring of reads, this is a non-strand-specific library. Orientation of splice sites (junction track) consistently show forward orientation on the left and reverse orientation on the right (red boxes), separated by one larger exon in the initial candidate transcript (red). Hence, a split is performed that results in two transcript models (green) that share a common exon and both contain a CDS prediction.

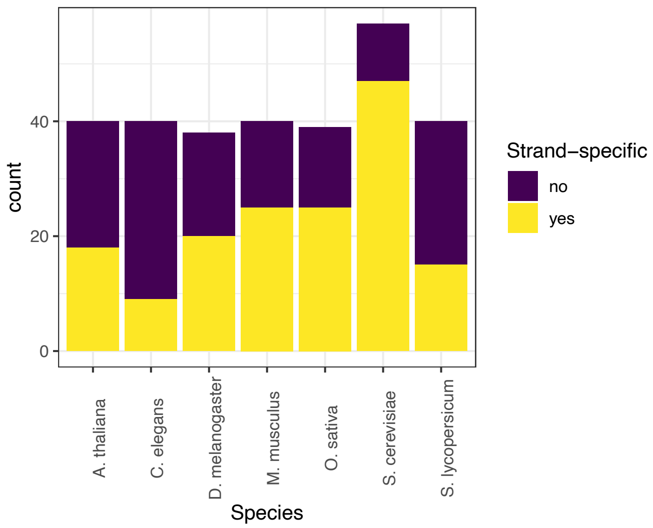


**Supplementary Figure S4.** Number of strand-specific and non-strand-specific libraries among the benchmark data sets.

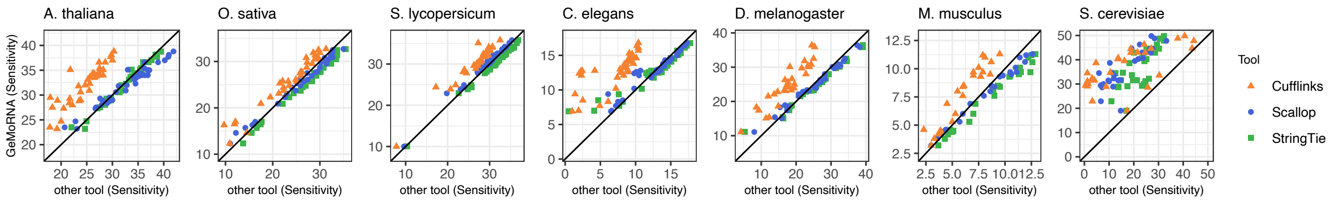


**Supplementary Figure S5.** Sensitivity according to gccompare. Benchmark results of GeMoRNA compared with Cufflinks, Scallop and StringTie based on the evaluation using gffcompare on the level of transcripts. Each panel displays the sensitivity of GeMoRNA (ordinate) compared with the remaining tools (abscissa, color coded). Points above the main diagonal indicate an improved performance of GeMoRNA, while points below the diagonal indicate an improved performance of the respective alternative tool.

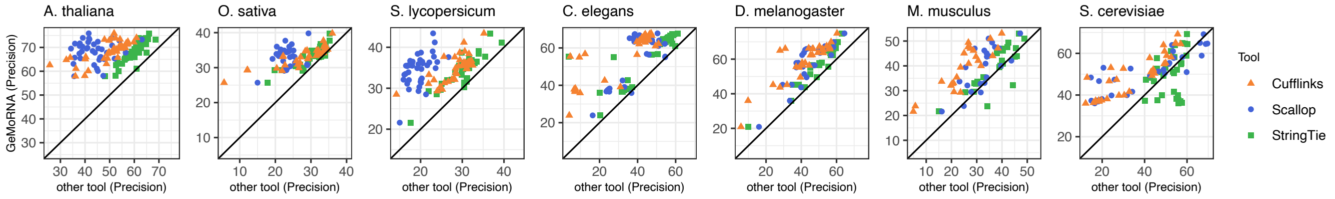


**Supplementary Figure S6.** Precision according to gffcompare. Benchmark results of GeMoRNA compared with Cufflinks, Scallop and StringTie based on the evaluation using gffcompare on the level of transcripts. Each panel displays the precision of GeMoRNA (ordinate) compared with the remaining tools (abscissa, color coded). Points above the main diagonal indicate an improved performance of GeMoRNA, while points below the diagonal indicate an improved performance of the respective alternative tool.

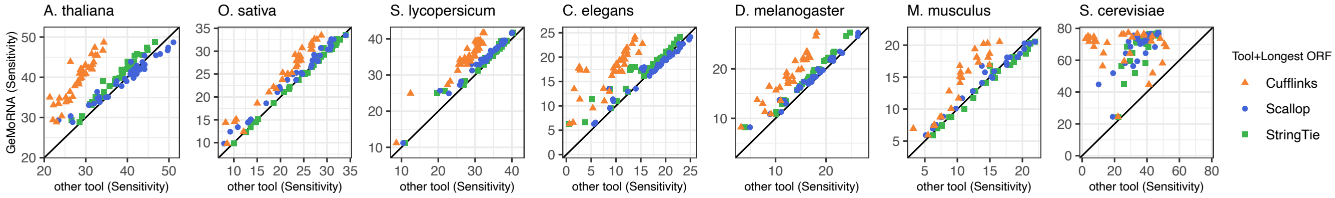


**Supplementary Figure S7.** Sensitivity according to GeMoMa Analyzer. Benchmark results of GeMoRNA compared with Cufflinks, Scallop and StringTie using the longest-ORF CDS prediction based on the evaluation using GeMoMa Analyzer on the level of CDS. Each panel displays the sensitivity of GeMoRNA (ordinate) compared with the remaining tools (abscissa, color coded). Points above the main diagonal indicate an improved performance of GeMoRNA, while points below the diagonal indicate an improved performance of the respective alternative tool.

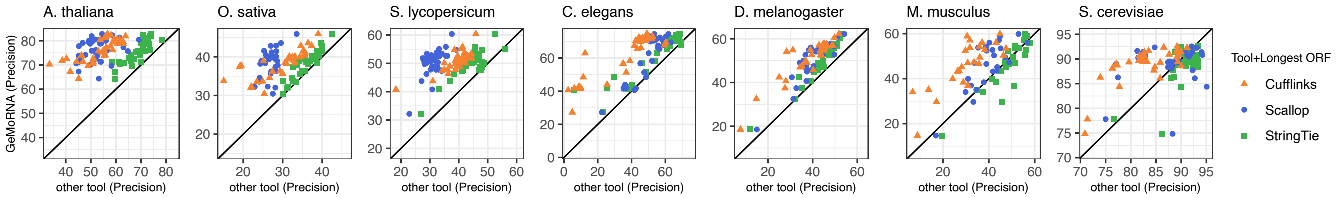


**Supplementary Figure S8.** Precision according to GeMoMa Analyzer. Benchmark results of GeMoRNA compared with Cufflinks, Scallop and StringTie using the longest-ORF CDS prediction based on the evaluation using GeMoMa Analyzer on the level of CDS. Each panel displays the precision of GeMoRNA (ordinate) compared with the remaining tools (abscissa, color coded). Points above the main diagonal indicate an improved performance of GeMoRNA, while points below the diagonal indicate an improved performance of the respective alternative tool.

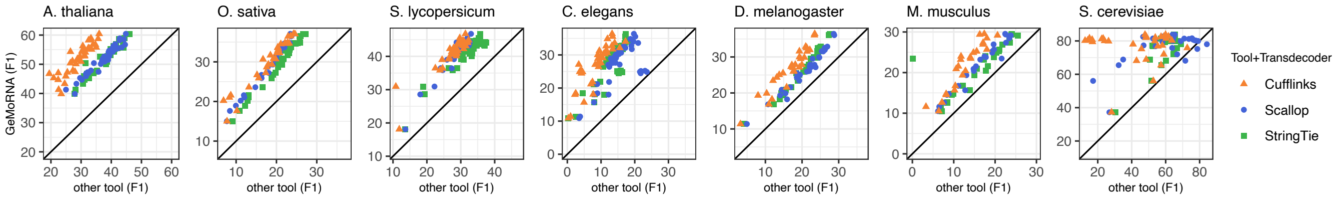


**Supplementary Figure S9.** F1 measure according to GeMoMa Analyzer. Benchmark results of GeMoRNA compared with Cufflinks, Scallop and StringTie using TransDecoder for CDS prediction based on the evaluation using GeMoMa Analyzer on the level of CDS. Each panel displays the sensitivity of GeMoRNA (ordinate) compared with the remaining tools (abscissa, color coded). Points above the main diagonal indicate an improved performance of GeMoRNA, while points below the diagonal indicate an improved performance of the respective alternative tool.

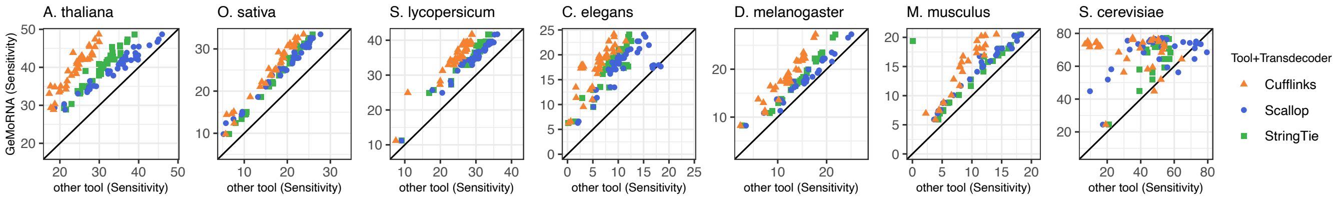


**Supplementary Figure S10.** Sensitivity according to GeMoMa Analyzer. Benchmark results of GeMoRNA compared with Cufflinks, Scallop and StringTie using TransDecoder for CDS prediction based on the evaluation using GeMoMa Analyzer on the level of CDS. Each panel displays the sensitivity of GeMoRNA (ordinate) compared with the remaining tools (abscissa, color coded). Points above the main diagonal indicate an improved performance of GeMoRNA, while points below the diagonal indicate an improved performance of the respective alternative tool.

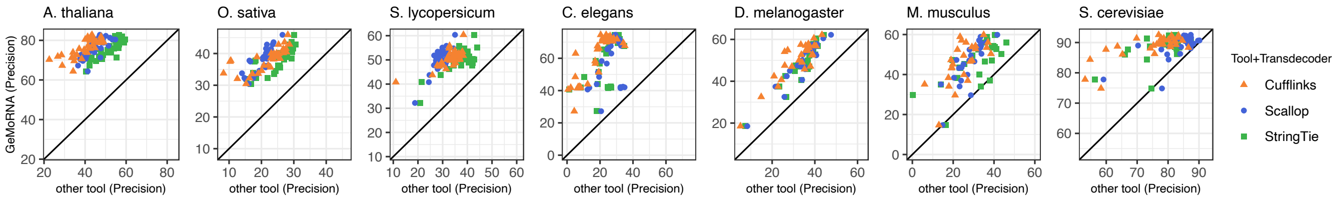


**Supplementary Figure S11.** Precision according to GeMoMa Analyzer. Benchmark results of GeMoRNA compared with Cufflinks, Scallop and StringTie using TransDecoder for CDS prediction based on the evaluation using GeMoMa Analyzer on the level of CDS. Each panel displays the precision of GeMoRNA (ordinate) compared with the remaining tools (abscissa, color coded). Points above the main diagonal indicate an improved performance of GeMoRNA, while points below the diagonal indicate an improved performance of the respective alternative tool.

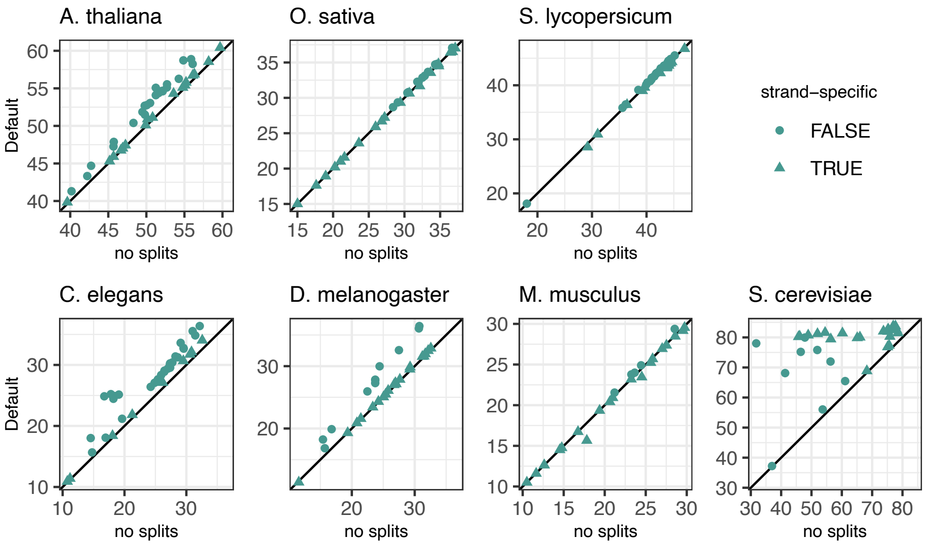


**Supplementary Figure S12.** Comparison of GeMoRNA with (ordinate) and without (abscissa) split heuristics using GeMoMa Analyzer on the level of CDS using the F1 measure. Points above the main diagonal indicate an improved performance when applying the split heuristics to candidate transcripts, while points below the diagonal indicate a loss in performance.

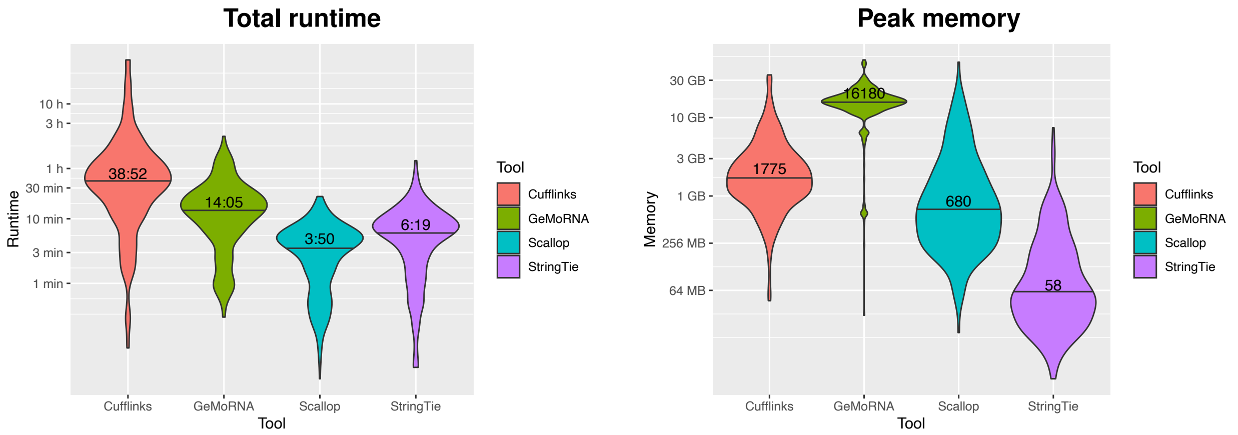


**Supplementary Figure S13.** Runtime and peak memory consumption of GeMoRNA, Cufflinks, Scallop and StringTie on the benchmark data sets.

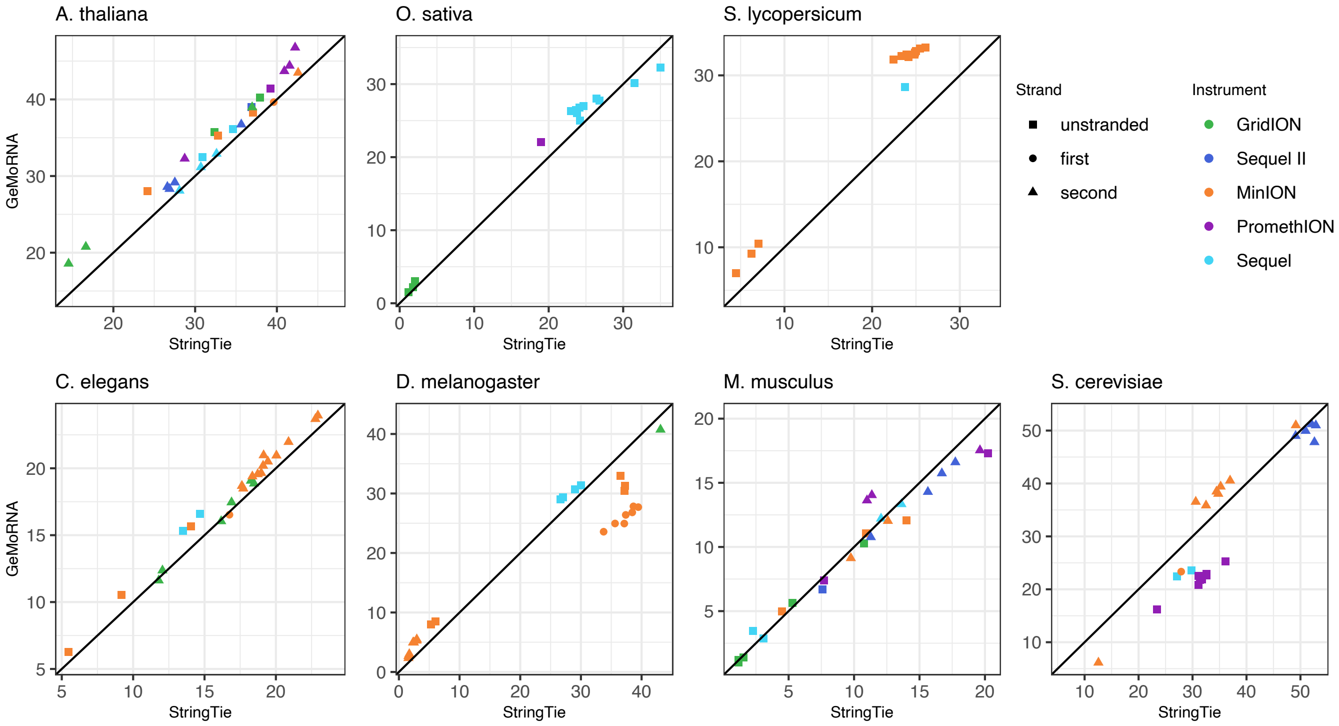


**Supplementary Figure S14.** F1 measure for long-read data on the level of transcripts according to gffcompare. Each panel displays the F1 measure of GeMoRNA (ordinate) compared with StringTie (abscissa). The instrument models that have been used for generating the respective long-read data are indicated by colour and strand specificity of indicated by shape. Points above the main diagonal indicate an improved performance of GeMoRNA, while points below the diagonal indicate an improved performance of StringTie.

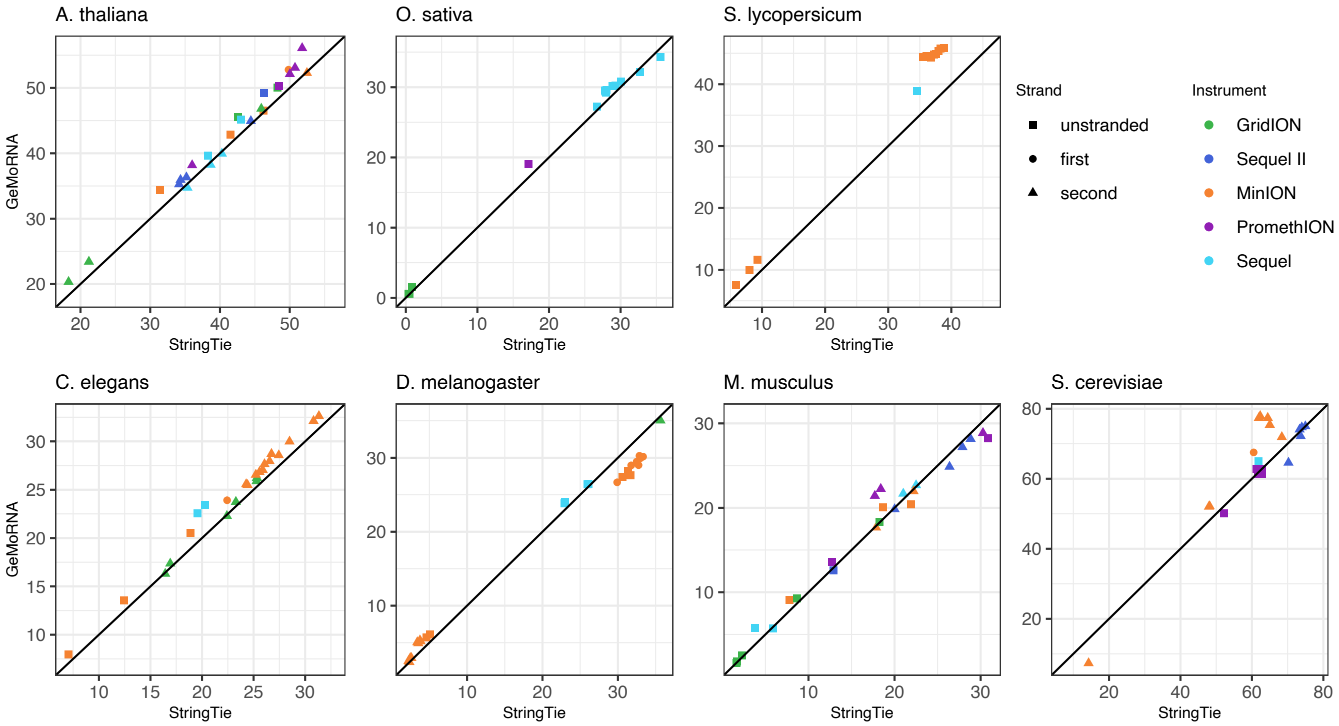


**Supplementary Figure S15.** F1 measure for long-read data on the level of CDS according to GeMoMa Analyzer. Each panel displays the F1 measure of GeMoRNA (ordinate) compared with StringTie using the longest-ORF CDS prediction (abscissa). The instrument models that have been used for generating the respective long-read data are indicated by colour and strand specificity of indicated by shape. Points above the main diagonal indicate an improved performance of GeMoRNA, while points below the diagonal indicate an improved performance of StringTie.

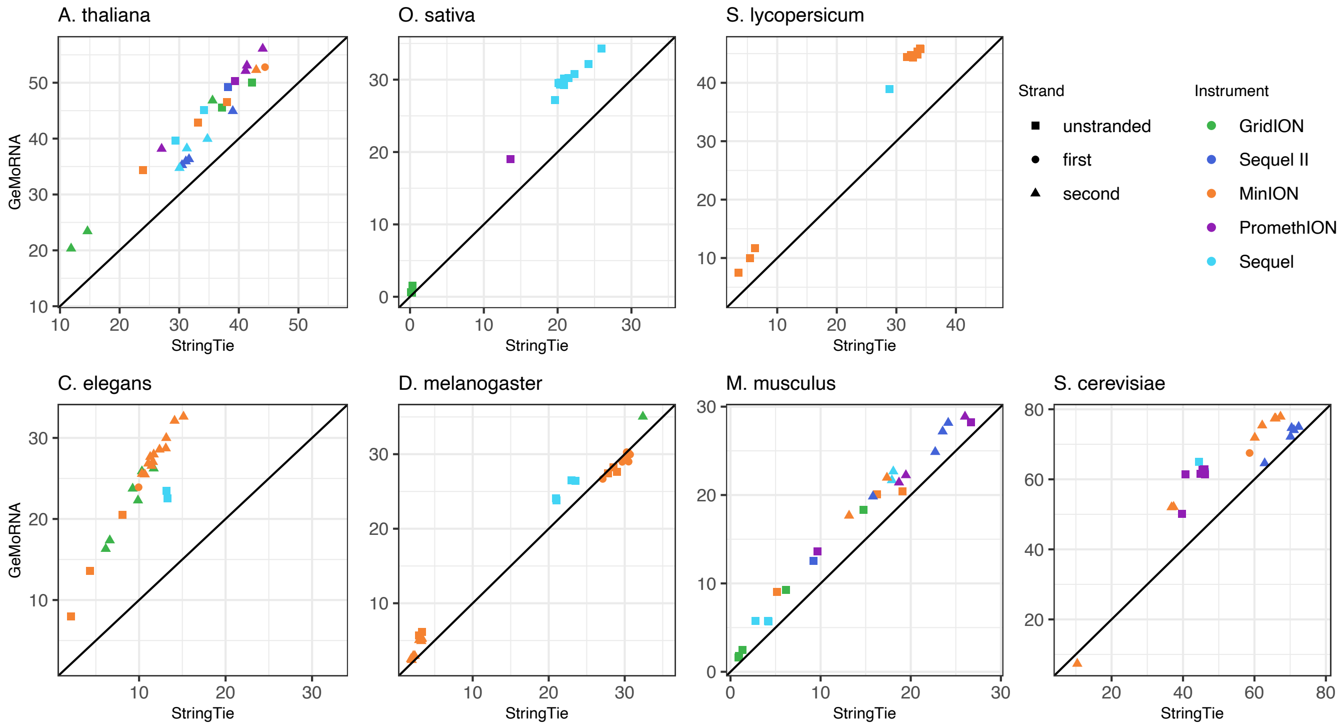


**Supplementary Figure S16.** F1 measure for long-read data on the level of CDS according to GeMoMa Analyzer. Each panel displays the F1 measure of GeMoRNA (ordinate) compared with StringTie using TransDecoder for CDS prediction (abscissa). The instrument models that have been used for generating the respective long-read data are indicated by colour and strand specificity of indicated by shape. Points above the main diagonal indicate an improved performance of GeMoRNA, while points below the diagonal indicate an improved performance of StringTie.

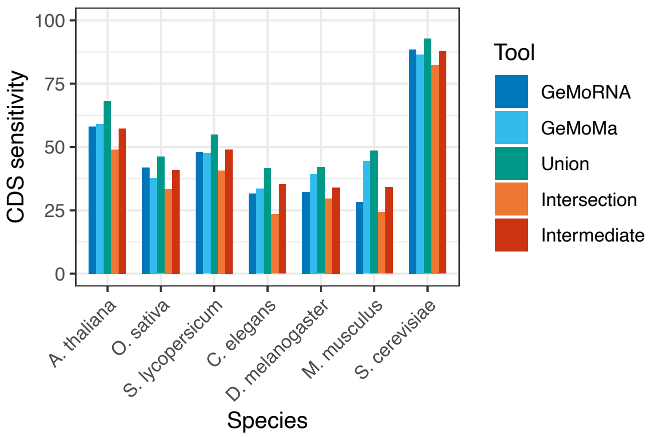


**Supplementary Figure S17.** Sensitivity on the level of CDS for GeMoRNA (RNA-seq-based) and GeMoMa (homology-based) predictions, and for combinations of the prediction sets.

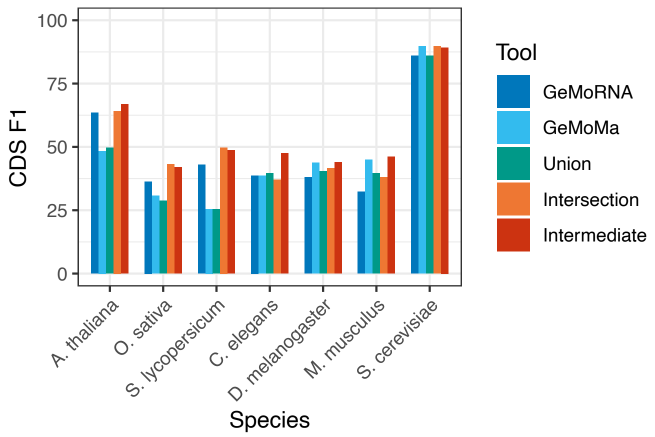


**Supplementary Figure S18.** F1 measure on the level of CDS for GeMoRNA (RNA-seq-based) and GeMoMa (homology-based) predictions, and for combinations of the prediction sets.

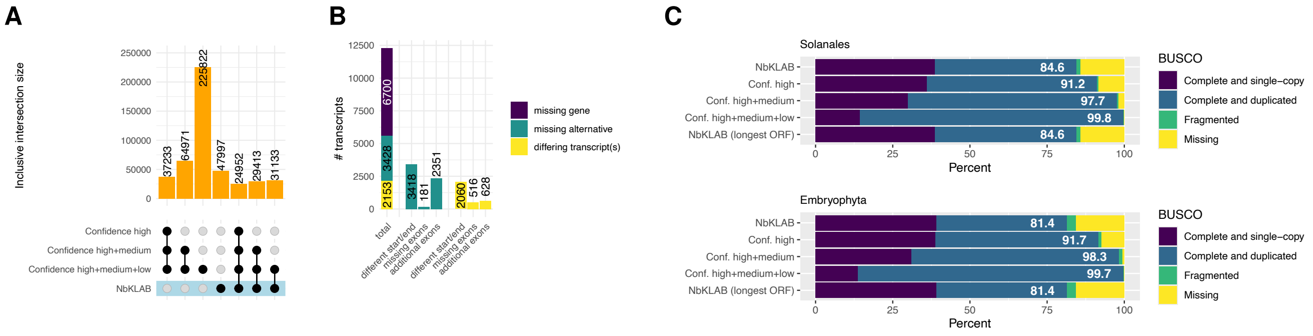


**Supplementary Figure S19.** Comparison of the original NbKLAB annotation to the merged prediction of GeMoRNA and GeMoMa considering transcripts of different levels of confidence. (A) Intersection sizes (inclusive) of the CDS contained in each of the annotation files. (B) Breakdown of transcripts that are present in the "Confidence high" set of GeMoRNA/GeMoMa but are missing from the NbKLAB annotation. (C) BUSCO completeness of the annotation files (extracted proteins) for the solanales and embryophyta datasets.

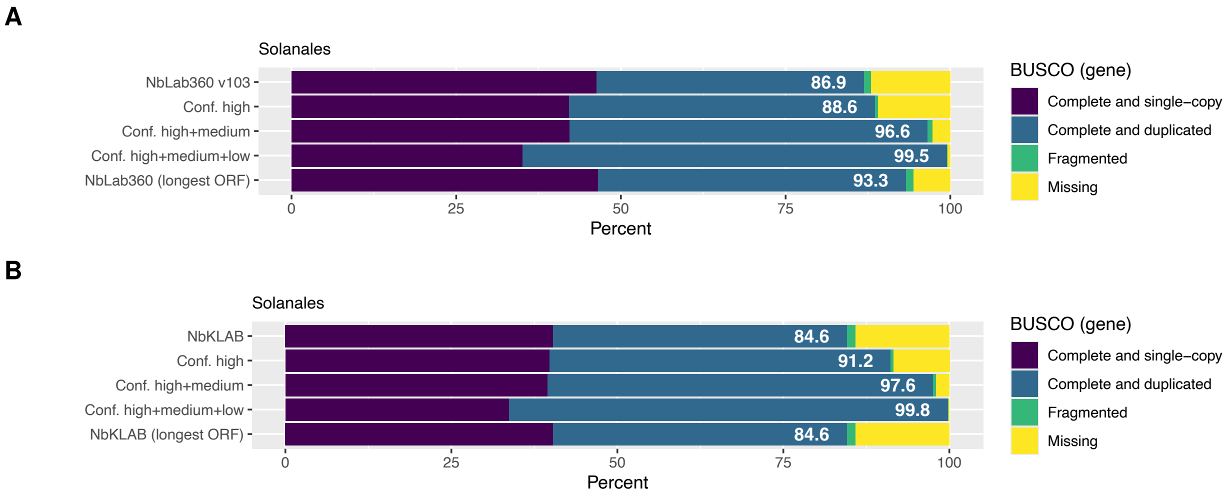


**Supplementary Figure S20.** BUSCO completeness (solanales, protein), determined on the level of genes using the BUSCORecomputer module of GeMoMa, of the annotation files for the NbLAB (A) and NbKLAB (B) genomes.

**A**

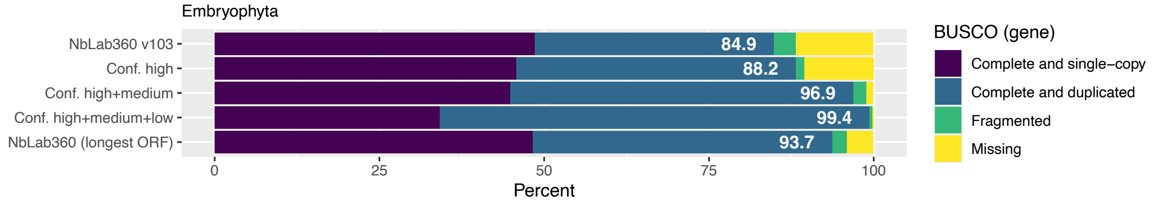


**B**

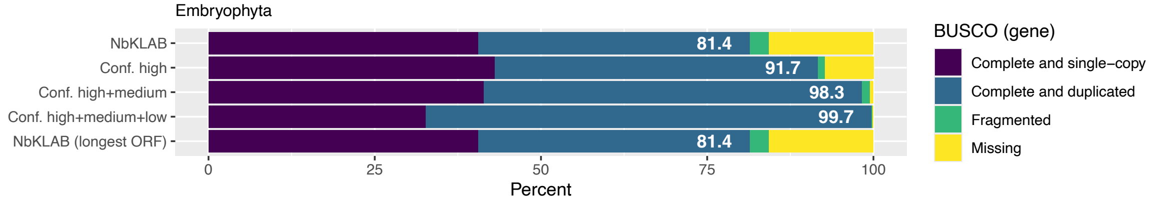


**Supplementary Figure S21.** BUSCO completeness (embryophyta, protein), determined on the level of genes using the BUSCORecomputer module of GeMoMa, of the annotation files for the NbLAB (A) and NbKLAB (B) genomes.
